# Fear, anxiety, and the extended amygdala—*Absence of evidence for strict functional segregation*

**DOI:** 10.1101/2025.08.29.673083

**Authors:** Paige R. Didier, Shannon E. Grogans, Claire M. Kaplan, Hyung Cho Kim, Samiha Islam, Allegra S. Anderson, Rachael M. Tillman, Manuel Kuhn, Juyoen Hur, Andrew S. Fox, Kathryn A. DeYoung, Jason F. Smith, Alexander J. Shackman

**Affiliations:** Department of Psychology, University of Maryland, College Park, MD 20742 USA; Neuroscience and Cognitive Science Program, University of Maryland, College Park, MD 20742 USA; Department of Maryland Neuroimaging Center, University of Maryland, College Park, MD 20742 USA; CommonSpirit Neuropsychology, Lakewood, CO 80228 USA; Rees-Jones Center for Foster Care Excellence, Children’s Heath, Dallas, TX 75207 USA; Department of Psychiatry and Human Behavior, Brown University, Providence, RI 02912 USA; McGill Neuropsychology, Bethesda, MD 20814 USA; Institute of Medical Psychology, Charité Universitätsmedizin Berlin, 10117 Berlin, Germany; Department of Psychology, Yonsei University, Seoul 03722, Republic of Korea; Department of Psychology, University of California, Davis, CA 95616 USA; California National Primate Research Center, University of California, Davis, CA 95616 USA

**Keywords:** affective neuroscience, fear and anxiety, bed nucleus of the stria terminalis (BST/BNST), extended amygdala (EA), Research Domain Criteria (RDoC)

## Abstract

Since the time of Freud, the distinction between fear and anxiety has been a hallmark of influential models of emotion and emotional illness, including the Diagnostic and Statistical Manual of Mental Disorders (DSM) and Research Domain Criteria (RDoC). Fear and anxiety disorders are common, debilitating, and challenging to treat, underscoring the importance of developing accurate models of the underlying neurobiology. Although there is consensus that the extended amygdala (EA) plays a central role in orchestrating responses to threat, the respective contributions of its two major subdivisions—the central nucleus of the amygdala (Ce) and bed nucleus of the stria terminalis (BST)—remain contentious. To help adjudicate this debate, we performed a harmonized mega-analysis of fMRI data acquired from 295 adults as they completed a well-established threat-anticipation paradigm. Contrary to popular double-dissociation models, results demonstrated that the Ce responds to temporally uncertain threat and the BST responds to certain threat. In direct comparisons, the two regions showed statistically indistinguishable responses, with strong Bayesian evidence of regional equivalence. In contrast, frontocortical regions responded preferentially to uncertain-threat anticipation. Together, these observations underscore the need to reformulate conceptual models that posit a strict segregation of temporally certain and uncertain threat processing in the EA.

## INTRODUCTION

Since the time of Freud, the distinction between fear and anxiety has been a hallmark of influential models of emotion and emotional illness, including the DSM and Research Domain Criteria (RDoC) (APA,2022; Freud et al., 1959; Grogans et al., 2023; Grupe & Nitschke, 2013; LeDoux & Pine, 2016; NIMH, 2011; Tovote et al., 2015). When extreme or pervasive, fear and anxiety can become debilitating (Salomon et al.,2015). Anxiety disorders are a leading cause of human misery, morbidity, and premature mortality (GBD2021, 2024). Existing treatments are far from curative for many, underscoring the need to develop a more complete and accurate understanding of the underlying neurobiology (Cuijpers et al., 2024; De Crescenzo et al., 2024; Singewald et al., 2023).

There is widespread consensus that the extended amygdala (EA)—a macrocircuit encompassing the central nucleus of the amygdala (Ce) and bed nucleus of the stria terminalis (BST)—plays a central role in fear and anxiety-related states, traits, and disorders, but the precise contributions of the Ce and BST remain contentious (Blanchard & Canteras, 2024; Daniel-Watanabe & Fletcher, 2022; Hur et al., 2020; Shackman et al., 2024). Building on an earlier generation of rodent studies (Shackman & Fox, 2016), RDoC and other double-dissociation models organize fear and anxiety into two strictly segregated neural systems (**Supplementary Figure S1 and Note 1**): the *Acute Threat* system is centered on the amygdala (including the Ce), is sensitive to certain (but not uncertain) threat, and promotes signs and symptoms of fear; whereas the *Potential Threat* system is centered on the BST, is sensitive to uncertain (but not certain) threat, and promotes anxiety (Avery et al., 2016; LeDoux & Pine, 2016; NIMH, 2011). Yet a growing body of rodent mechanistic data casts doubt on this strict either/or perspective (Ahrens et al.,2018; Bruzsik et al., 2021; Chen et al., 2022; Griessner et al., 2021; Gungor & Paré, 2016; Holley & Fox, 2022; Lange et al., 2017; Lee et al., 2017; Marcinkiewcz et al., 2016; Marvar et al., 2021; Moscarello & Penzo, 2022; Pomrenze, Giovanetti, et al., 2019; Pomrenze, Tovar-Diaz, et al., 2019; Ren et al., 2022; Ressler et al., 2020; Zhu et al., 2024), motivating the competing hypothesis that the Ce and BST play a role in organizing responses to both kinds of threat (Daniel-Watanabe & Fletcher, 2022; Fox & Shackman,2019; Gungor & Paré, 2016).

To help adjudicate this debate, we performed a harmonized mega-analysis of fMRI data acquired from 295 racially diverse adults as they completed the Maryland Threat Countdown (MTC), a well-established threat-anticipation paradigm (**Figure 1**) (Hur et al., 2020). The MTC is an fMRI-optimized variant of temporally certain/uncertain-threat assays that have been behaviorally and pharmacologically validated in rodents and humans, maximizing translational relevance (Hur et al., 2020). Data were acquired using a multiband sequence and re-processed using a singular best-practices pipeline. The relatively large sample afforded the power necessary to reliably detect small differences in regional responses to certain- and uncertain-threat anticipation (*d*≥0.16).

**Figure 1.**
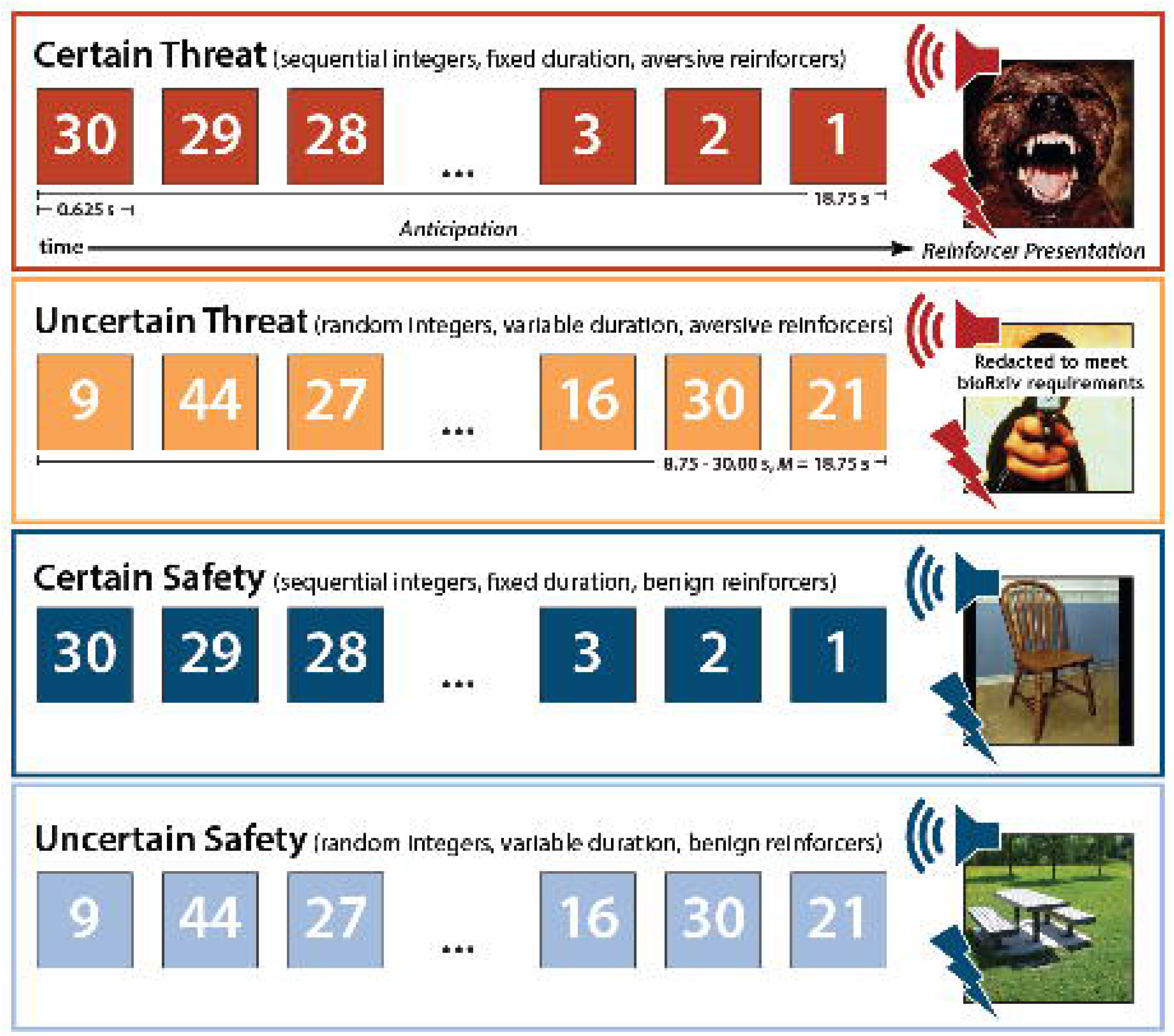
Maryland Threat Countdown fMRI Paradigm. The paradigm takes the form of a 2 (*Valence:* Threat, Safety) × 2 (*Temporal Certainty:* Certain, Uncertain) randomized event-related design. Participants were completely informed about the task design and contingencies prior to scanning. On certain-threat trials, participants saw a descending stream of integers (‘countdown’) for 18.75 s. To ensure robust distress and arousal, the anticipation epoch always terminated with the presentation of a noxious electric shock, unpleasant photograph, and thematically related audio clip (e.g., scream). Uncertain-threat trials were similar, but the integer stream was randomized and presented for an uncertain and variable duration (8.75–30.00 s; *M*=18.75 s). Participants knew that something aversive was going to occur, but they had no way of knowing precisely *when*. Safety trials were similar but terminated with the delivery of emotionally neutral reinforcers (e.g., just-perceptible electrical stimulation). Abbreviation—s, seconds.

Because voxelwise analyses do not permit inferences about regional differences, we used *a priori* anatomical regions of interest (ROIs) to rigorously compare the Ce and BST. This has the advantage of providing statistically unbiased effect-size estimates (Poldrack, Baker, Durnez, Gorgolewski, Matthews, Munafò, et al., 2017), in contrast to earlier work by our group that relied on functionally defined ROIs (Hur et al., 2020). To maximize anatomical resolution and inferential clarity, mean activation was computed using spatially unsmoothed data. Hypothesis testing focused on regional responses to certain- and uncertain-threat anticipation relative to their perceptually similar reference conditions (e.g., certain-safety anticipation), providing sharper inferences than prior work focused on baseline contrasts (Grogans et al., 2024). Of course, traditional null-hypothesis tests cannot address whether the Ce and BST show statistically equivalent responses to certain- and uncertain-threat anticipation. Here we used a Bayesian framework to quantify the strength of the evidence for and against regional differences. The Bayesian approach provides well-established benchmarks for interpreting effect sizes and sidesteps the need to arbitrarily choose what constitutes a ‘statistically indistinguishable’ difference (Bo et al., 2024), unlike work focused on frequentist equivalence tests (Shackman et al., 2024). Whole-brain voxelwise analyses enabled us to explore less intensively scrutinized regions.

## METHOD

### Overview of the Mega-Analysis

The neuroimaging mega-analysis capitalized on data from two previously published fMRI studies focused on the neural circuits recruited by temporally certain- and uncertain-threat anticipation. The first study encompassed a sample of 220 psychiatrically healthy, first-year university students (Grogans et al., 2024). The second encompassed 75 tobacco smokers recruited from the surrounding community (Kim et al., 2023). Both studies employed the same certain/uncertain threat-anticipation paradigm (Maryland Threat Countdown task) and were collected using identical parameters on the same scanner using the same head-coil. For the mega-analysis, all neuroimaging data were completely reprocessed using a singular best-practices pipeline, as detailed below. This harmonized approach is increasingly common and generally provides greater statistical power and generalizability, relative to typical single-study approaches (Petre et al., 2022; Radua et al., 2025), and greater anatomical resolution and statistical rigor, as compared to coordinate-based meta-analytic approaches (Costafreda, 2009; Salimi-Khorshidi et al., 2009). All participants provided informed written consent. Procedures were approved by the University of Maryland, College Park Institutional Review Board (protocols #659385 and #824438).

Detailed descriptions of the study designs, enrollment criteria, participants, data collection procedures, and data exclusions are provided in the original reports (Grogans et al., 2024; Kim et al., 2023). The mega-analysis was not pre-registered.

### Participants

Across studies, a racially diverse sample of 295 participants provided usable neuroimaging data (Total: 45.4% female; 52.2% White Non-Hispanic, 16.6% Asian, 19.0% African American, 4.1% Hispanic, 8.1% Multiracial/Other; *M*=21.6 years, *SD*=5.7, *Range*=18-40 years; University Students: 49.5% female; 61.4% White Non-Hispanic, 18.2% Asian, 8.6% African American, 4.1% Hispanic, 7.3% Multiracial/Other; *M*=18.8 years; *SD*=0.4; Community Tobacco Smokers: 33.3% female; 25.3% White Non-Hispanic, 12.0% Asian, 49.3% African American, 4.0% Hispanic, 9.4% Multiracial/Other; *M*=30.1 years; *SD*=5.6). Of these, 8 participants were excluded from skin conductance analyses due to insufficient usable data (for additional details regarding data censoring, see Grogans et al., 2024; Kim et al., 2023).

### Power Analysis

To enable readers to better interpret in our results, we performed a post hoc power analysis. G-power (version 3.1.9.2) indicated that the final sample of 295 usable fMRI datasets provides 80% power to detect ‘small’ mean differences in regional activation (Cohen’s *d*=0.16, α=0.05, two-tailed) (Cohen, 1988; Faul et al., 2007).

### Threat-Anticipation Paradigm

As shown in **Figure 1**, the Maryland Threat Countdown paradigm takes the form of a 2 (*Valence:* Threat/Safety) × 2 (*Temporal Certainty:* Uncertain/Certain) design. On Certain-Threat trials, participants saw a descending stream of integers (‘count-down’) for 18.75 s. To ensure robust emotion, the anticipation epoch culminated with the presentation of a noxious electric shock, unpleasant photograph, and thematically related audio clip. Uncertain-Threat trials were similar, but the integer stream was randomized and presented for an uncertain and variable duration (8.75-30.00 s; *M*=18.75 s). Participants knew that something aversive was going to occur but had no way of knowing precisely when. The Maryland paradigm differs from threat-anticipation tasks (e.g., NPU) that manipulate the probability (or probability *and* timing) of encounters (Schmitz & Grillon, 2012). For additional details, see the **Supplement**. Participants were periodically prompted to rate the intensity of fear/anxiety experienced a few seconds earlier, during the anticipation period of the prior trial, using a 1 (*minimal*) to 4 (*maximal*) scale. Skin conductance was continuously acquired throughout (for details, see the **Supplement**).

### MRI Data Acquisition

Data were acquired using a Siemens Magnetom TIM Trio 3 Tesla scanner (32-channel head-coil; for additional details, see the **Supplement**). T1-weighted anatomical images were acquired using a magnetization prepared rapid acquisition gradient echo sequence (TR=2,400 ms; TE=2.01 ms; inversion time=1,060 ms; flip=8°; slice thickness=0.8 mm; in-plane=0.8×0.8 mm; matrix=300×320; field-of-view=240×256). A T2-weighted image was collected co-planar to the T1-weighted image (TR=3,200 ms; TE=564 ms; flip angle=120°). A multi-band sequence was used to collect oblique-axial echo-planar imaging (EPI) volumes (acceleration=6; TR=1,250 ms; TE=39.4 ms; flip=36.4°; slice thickness=2.2 mm, slices=60; in-plane =2.1875×2.1875 mm; matrix=96×96; 3×478-volume scans). Co-planar oblique-axial spin echo (SE) images were collected in opposing phase-encoding directions (rostral-to-caudal and caudal-to-rostral; TR=7,220 ms; TE=73 ms). Respiration and pulse were continuously acquired.

### MRI Pipeline

Methods are similar to other work and are only briefly summarized here (Cornwell et al., 2025). For details, see the **Supplement**.

### Anatomical Data Processing

T1- and T2-weighted images were inhomogeneity corrected, denoised, and brain-extracted. Extracted T1 images were diffeomorphically normalized to a 1-mm T1-weighted MNI152 template using *ANTS*.

### Functional Data Processing

EPI files were de-spiked, slice-time corrected, motion-corrected, inhomogeneity corrected, co-registered using boundary-based registration, normalized to the MNI template, and resampled (2 mm^3^). Voxelwise analyses employed data that were spatially smoothed (4-mm). Extended amygdala (EA) analyses used spatially unsmoothed data and anatomical regions-of-interest (ROIs; see below).

### fMRI Data Modeling

Methods are similar to other recent work and only briefly summarized here (Cornwell et al., 2025). For additional details, see the **Supplement**.

#### First-Level Modeling

Modeling was performed using *SPM12*, the default autoregressive model, and a temporal band-pass filter set to the hemodynamic response function (HRF) and 128 s set to the hemodynamic response function (HRF) and 128 s (0.0078-0.1667 Hz). Regressors were convolved with a canonical HRF and temporal derivative. For the threat-anticipation paradigm, hemodynamic activity was modeled using rectangular regressors spanning the entirety of the anticipation (‘countdown’) epochs of the Uncertain-Threat, Certain-Threat, and Uncertain-Safety trials. Certain-Safety anticipation served as the reference condition and contributed to the implicit baseline estimate. Epochs corresponding to the reinforcers, visual masks, and rating prompts were modeled using the same approach. Nuisance variates included volume-to-volume displacement and first derivative, 6 motion parameters and first derivatives, cerebrospinal fluid (CSF), pulse, respiration, and other nuisance signals (e.g., brain edge, CSF edge, global motion, WM, and extracerebral soft tissue). Volumes with excessive displacement (>0.5 mm) and those during and immediately following reinforcer (shock) delivery were censored.

#### ROIs

Ce and BST activation was quantified using anatomical ROIs, unsmoothed data, and regression coefficients extracted and averaged for each combination of contrast, region, and participant (Theiss et al., 2017; Tillman et al., 2018). Anatomical ROIs enable statistically unbiased tests of regional sensitivity to specific experimental manipulations (i.e., Region × Condition effects), including potential single and double dissociations (e.g., BST: Uncertain > Certain Threat; Ce: Uncertain < Certain Threat) (Fox et al., 2018; Poldrack, Baker, Durnez, Gorgolewski, Matthews, Munafo, et al., 2017). ROI registration appeared reasonable for both regions and hemispheres, based on visual inspection of the 295 normalized T1-weighted anatomical images.

### Analytic Strategy

Methods are similar to other work and are only briefly summarized here (Cornwell et al., 2025). For details, see the **Supplement**.

#### Overview

Frequentist (Cohen’s *d*) effect sizes were interpreted using established benchmarks (Cohen, 1988; Cohen, 1994; Schimmack, 2019), ranging from *large* (*d*=0.80), to *medium* (*d*=0.50), to *small* (*d*=0.20), to *nil* (*d*≤0.10). Bayesian effect sizes were computed for select analyses. Bayes Factor (BF^10^) quantifies the relative performance of the null hypothesis (*H*_*0*_; e.g., the absence of a credible mean difference) and the alternative hypothesis (*H*_*1*_; e.g., the presence of a credible mean difference), on a 0 to ∞ scale. BF can be used to quantify the relative strength of the evidence for *H*_*0*_ (test the null), unlike conventional frequentist null-hypothesis significance tests (Bo et al., 2024; Wagenmakers et al., 2018). It also does not require the data analyst to arbitrarily decide what constitutes a ‘statistically indistinguishable’ difference, in contrast to traditional equivalence tests (Hur et al., 2020). This approach provides readily interpretable, principled effect-size benchmarks (van Doorn et al., 2021). Values >1 were interpreted as evidence of mean differences in activation across conditions, ranging from *strong* (*BF*_*10*_>10), to *moderate* (*BF*_*10*_=3-10), to *weak* (*BF*_*10*_=1-3). Values <1 were interpreted as evidence of statistical equivalence (i.e., support for the null hypothesis), ranging from s*trong* (*BF*_*10*_<0.10), to *moderate* (*BF*_*10*_=0.10-0.33), to *weak* (*BF*_*10*_=0.33-1). The reciprocal of *BF*_*10*_ represents the relative likelihood of the null hypothesis (e.g., *BF*_*10*_=0.10, *H*_*0*_ is 10 times more likely than *H*_*1*_). Bayesian effects were computed using a noninformative zero-centered Cauchy distribution 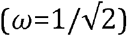, the field standard for two-sided tests (Gronau et al., 2020; Schmalz et al., 2023; Schönbrodt et al., 2017; van Doorn et al., 2021; Wagenmakers et al., 2018). Across tests, the estimated error of the MCMC-derived (Markov Chain Monte Carlo) *BF*_*10*_ estimates was negligible (<0.30%) and stable across a range of priors.

#### Whole-Brain Voxelwise Tests

Whole-brain voxelwise repeated-measures GLMs were used to compare each threat-anticipation condition to its corresponding control condition (e.g., Uncertain-Threat vs. Uncertain-Safety anticipation), while accounting for mean-centered study (Grogans et al., 2024; Kim et al.,2023), age, and assigned sex. A minimum conjunction was used to identify voxels sensitive to Certain-*and* Uncertain-Threat anticipation (Nichols et al., 2005). We also directly compared Certain-to Uncertain-Threat anticipation.

#### ROIs

One-sample Student’s *t*-tests was used to confirm that the EA ROIs showed nominally significant recruitment during Certain and Uncertain Threat anticipation (*p*<0.05, uncorrected). We used a standard 2 (*Region:* Ce, BST) × 2 (*Threat*-*Certainty:* Certain, Uncertain) repeated-measures GLM to test regional differences in activation during the anticipation of temporally Certain Threat (relative to Certain Safety) versus Uncertain Threat (relative to Uncertain Safety). Interactions were probed using focal contrasts. Sensitivity analyses confirmed that none of the conclusions materially changed when controlling for study, age, and assigned sex (for details, see the study OSF collection). A sign test (*Z*_*Sign*_) was used to test the proportion of participants showing double dissociations.

## RESULTS

### Threat anticipation amplifies subjective distress and objective arousal

We used repeated-measures general linear models (GLMs) to confirm that the threat-anticipation paradigm had the intended impact on anticipatory distress (in-scanner ratings) and arousal (skin conductance level, SCL). Eight participants were excluded from SCL analyses due to insufficient usable data (*n*=287). As shown in **Figure 2A**, subjective feelings of fear and anxiety were significantly elevated during the anticipation of threat compared to safety, and distress was particularly pronounced when the timing of threat encounters was uncertain (*Valence: F*(1,294)=965.74, *p*<0.001, *d*=1.81 [1.62, 1.99], *BF*_*10*_=1.42×10^91^; *Certainty: F*(1,294)=231.95, *p*<0.001, *d*=0.89 [0.75, 1.02], *BF*_*10*_=3.91×10^35^; *Valence × Certainty: F*(1,294)=25.58, *p*<0.001, *d*=0.29 [0.18, 0.41], *BF*_*10*_=12,327.09; *Threat, Uncertain vs. Certain: F*(1,294)=154.04, *p*<0.001, *d*=0.72 [0.59, 0.85], *BF*_*10*_=3.14×10^25^; *Safety, Uncertain vs. Certain: F*(1,218)=77.63, *p*<0.001, *d*=0.51 [0.39, 0.63], *BF*_*10*_=4.42×10^13^).

**Figure 2.**
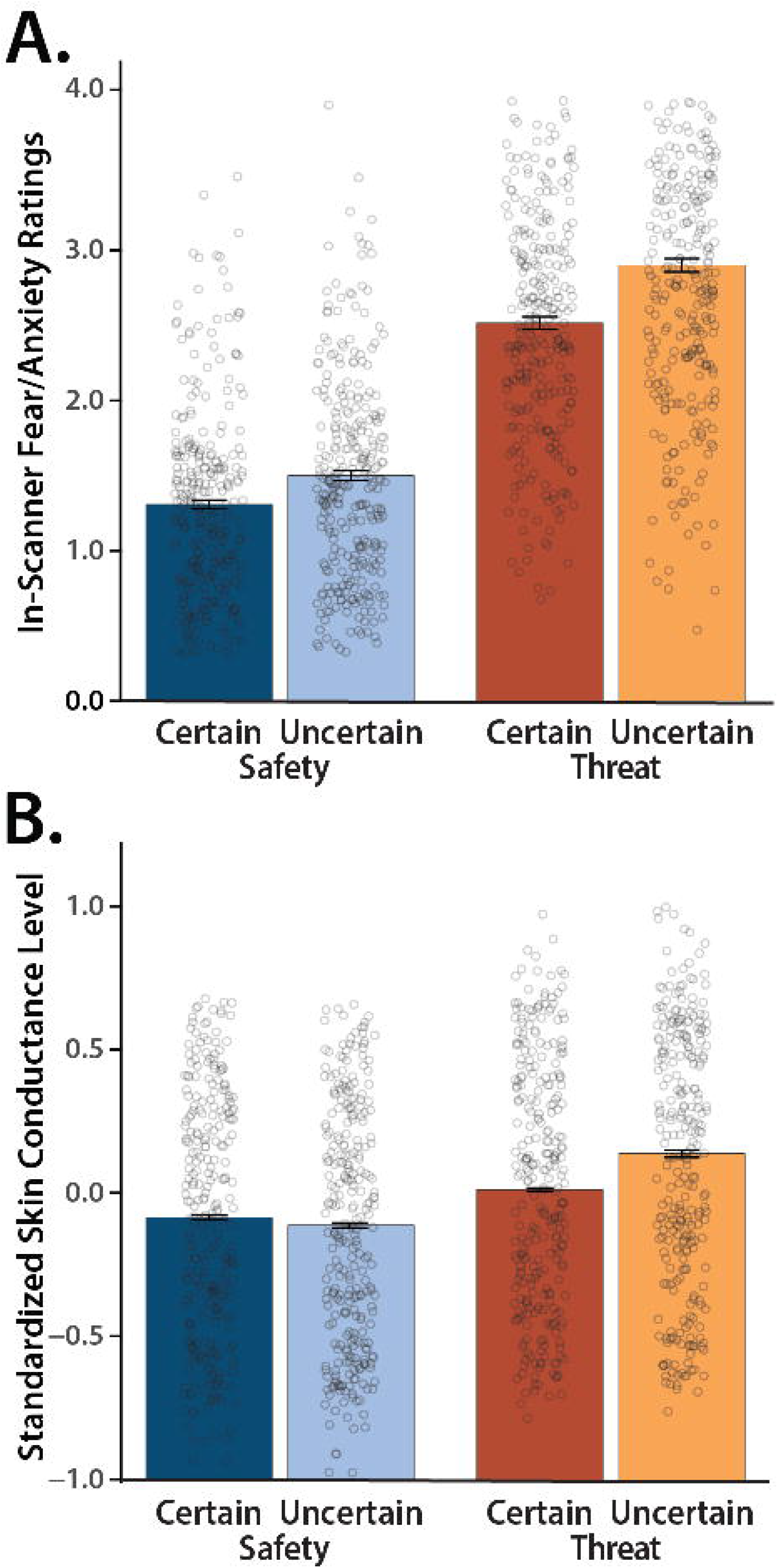
The Maryland Threat Countdown paradigm is a valid probe of human fear and anxiety. (A) Anticipated threat increases subjective symptoms of distress. Conscious feelings of fear and anxiety were increased during the anticipation of threat compared to safety, and this was particularly evident for temporally uncertain threat (*p*<0.001). **(B) Anticipated threat increases objective signs of arousal**. A similar pattern was evident for skin conductance level (SCL; *p*<0.001). Bars depict means, whiskers depict standard errors, and open rings depict individual participants.

As shown in **Figure 2B**, the same general pattern was evident for SCL, an objective psychophysiological index of anticipatory arousal (*Valence: F*(1,286)=165.76, *p*<0.001, *d*=0.76 [0.63, 0.89], *BF*_*10*_=9.61×10^26^; *Certainty: F*(1,286)=80.21, *p*<0.001, *d*=0.53 [0.41, 0.65], *BF*_*10*_=1.09×10^14^; *Valence × Certainty: F*(1,286)=129.87, *p*<0.001, *d*=0.67 [0.54, 0.80], *BF*_*10*_=7.53×10^21^; *Threat, Uncertain vs. Certain: F*(1,286)=120.97, *p*<0.001, *d*=0.65 [0.52, 0.78], *BF*_*10*_=3.49×10^20^; *Safety, Uncertain vs. Certain: F*(1,286)=43.61, *p*<0.001, *d*=-0.39 [-0.51, -0.27], *BF*_*10*_=3.57×10^7^). Taken together, these converging observations confirm the validity of the MTC paradigm as an experimental probe of human fear and anxiety, consistent with work in smaller samples (Hur et al., 2020; Kim et al., 2023).

### Uncertain-threat anticipation recruits a distributed cortico-subcortical network

We used a whole-brain voxelwise GLM to identify regions recruited during the anticipation of temporally uncertain threat, relative to uncertain safety (*p*<0.05, whole-brain FWE corrected). As shown in the first column of **Figure 3**, this revealed a widely distributed network of cortical and subcortical regions previously implicated in the expression and regulation of human fear and anxiety (Bo et al., 2024; Chavanne & Robinson, 2021; Grogans et al., 2024; Hur et al., 2022; Radua et al., 2025; Shackman & Fox, 2021), including the midcingulate cortex (MCC); anterior insula (AI) extending into the frontal operculum (FrO); dorsolateral prefrontal cortex (dlPFC) extending to the frontal pole (FP); brainstem encompassing the periaqueductal grey (PAG); basal forebrain, including the BST; and dorsal amygdala, including the Ce (**Supplementary Table S1**).

**Figure 3.**
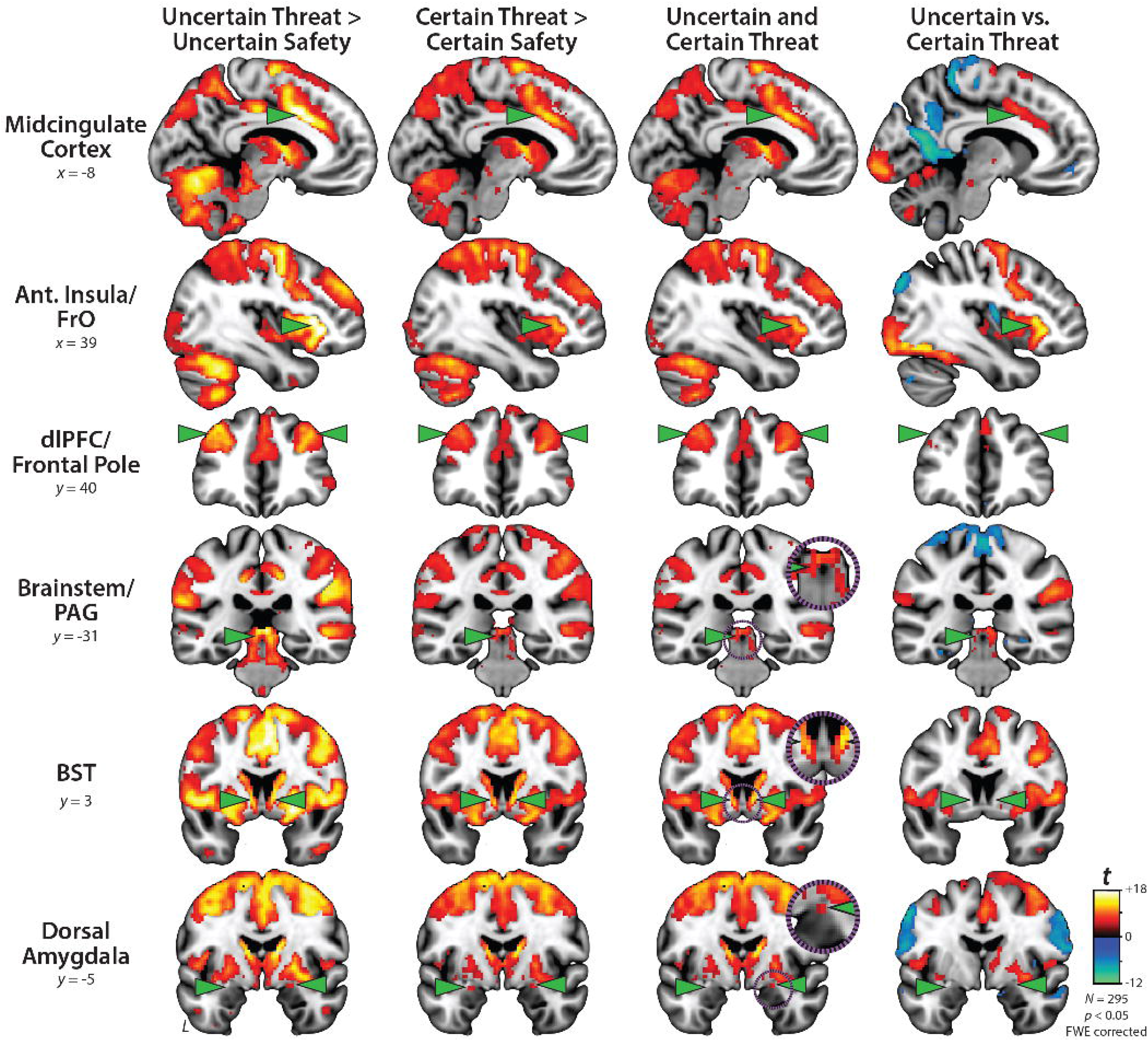
Uncertain- and certain-threat anticipation recruit a common cortico-subcortical network. Key regions (*green arrowheads*) show significantly increased activation during the anticipation of both uncertain threat (*first column*) and certain threat (*second column*), relative to their respective control conditions (*p*<0.05, whole-brain FWE corrected). *Third column* depicts the voxelwise conjunction (logical ‘AND’) of the two thresholded contrasts. Co-localization is evident throughout the network, including the BST and dorsal amygdala (Ce). *Fourth column* shows the direct contrast of the two threat-anticipation conditions. The MCC, AI/FrO, and to a lesser extent dlPFC/FP, show significantly greater activation during the anticipation of uncertain threat, whereas the BST, dorsal amygdala (Ce), and PAG show negligible discrimination of the two conditions. The dlPFC/FP mean difference was more evident at more rostral planes. For additional details, see **Supplementary Tables S1-S6**. *Purple insets* depict magnified views of overlap in the PAG, BST, and Ce. Abbreviations—Ant., Anterior; BST, Bed Nucleus of the Stria Terminalis; dlPFC, Dorsolateral Prefrontal Cortex; FrO, Frontal Operculum; FWE, Familywise Error; PAG, Periaqueductal Gray.

While not the focus of our study, exploratory analyses indicated that uncertain-threat anticipation was associated with reduced activation (‘de-activation’) in a set of midline regions that encompassed key nodes of the default mode network (e.g., frontal pole, rostral and straight gyri, and precuneus) as well as the pre- and post-central gyri, posterior insula, parahippocampal gyrus, and hippocampus (**Supplementary Table S2**), dovetailing with prior observations (Choi et al., 2012; Grupe et al., 2016; Hur et al., 2020). At a more liberal threshold (FDR *q*<0.05), the same pattern was evident in ventromedial regions of the amygdala (e.g., basal and cortical nuclei and amygdalohippocampal transition area), consistent with prior neuroimaging studies of anticipated threat and with the known functional heterogeneity of this complex structure (Cornwell et al., 2025; Fox & Shackman, 2024; Murty et al., 2022; Murty et al., 2023).

### Certain-threat anticipation recruits a broadly similar network

We used a parallel approach to identify regions recruited during the anticipation of temporally certain threat, relative to certain safety (*p*<0.05, whole-brain FWE corrected). As shown in the second column of **Figure 3**, the results strongly overlapped those evident for uncertain threat (**Supplementary Tables S3-S4**). Indeed, a minimum-conjunction analysis of the two contrasts (Logical ‘AND;’ Nichols et al., 2005) revealed voxelwise colocalization in every key region, including the BST and dorsal amygdala in the region of the Ce (third column of **Figure 3**). Taken together, these results suggest that this distributed cortico-subcortical system is sensitive to a range of anticipated threats, including those that are certain and uncertain in their timing.

### Frontocortical regions discriminate uncertain from certain threat, subcortical regions do not

To determine whether regions recruited during threat anticipation are sensitive to temporal uncertainty, we directly compared the uncertain and certain threat conditions (*p*<0.05, whole-brain FWE corrected). As shown in the fourth column of **Figure 3**, frontocortical regions—including MCC, AI/FrO, and dlPFC/FP—while engaged by both kinds of threat, showed a preference for temporally uncertain threat, consistent with prior work (**Supplementary Tables S5-S6**) (Hur et al., 2020). In contrast, the BST, dorsal amygdala (Ce), and PAG showed negligible differences.

### The BST and Ce show statistically indistinguishable responses to certain- and uncertain-threat anticipation

Because voxelwise analyses do not permit inferences about regional differences in activation, we used anatomical regions of interest (ROIs) and spatially unsmoothed data to rigorously compare the BST and Ce (**Figure 4A**). As a precursor to hypothesis testing, we used one-sample *t*-tests to confirm that the BST and Ce ROIs show significant activation during certain- and uncertain-threat anticipation relative to their respective control conditions (*t*(294)>6.49, *p*<0.001, *d*>0.37, *BF*_*10*_>2.02 × 10^7^). Next, we used a standard 2 (*Region:* BST, Ce) × 2 (*Threat*-*Certainty:* Certain, Uncertain) repeated-measures GLM to probe potential regional differences in threat sensitivity. Here again the BST and Ce proved statistically indistinguishable (**Figure 4A, Supplementary Figure S2A-B**). The critical Region × Threat-Certainty contrast was not significant (*F*(1,294)=0.12, *p*=0.73, *d*=-0.02 [-0.13, 0.09], *BF*_*10*_=0.07). In fact, participants were just as likely as not (49.5% vs. 50.5%; *H*_*0*_=50.0%) to show the hypothesized double-dissociation pattern (*Z*_*Sign*_=0.12, *p*=0.91; **Figures 4B, Supplementary Figure S2C**). Focal contrasts indicated that neither the BST nor the Ce credibly discriminated certain- from uncertain-threat anticipation (*BST: t*(294)=-0.59, *p*=0.56, *d*=-0.03 [-0.15, 0.08], *BF*_*10*_=0.11; *Ce: t*(294)=-1.38, *p*=0.17, *d*=-0.08 [-0.19, 0.03], *BF*_*10*_=0.03; **Figure 4A**), consistent with the more conservatively thresholded voxelwise results (**Figure 3**). The GLM did, however, reveal a main effect of region, reflecting generally greater BST reactivity to both kinds of anticipated threat (*F*(1,294)=95.36, *p*<0.001, *d*=0.57 [0.45, 0.69], *BF*_*10*_=3.95 × 10^16^). The main effect of Threat-Certainty was not significant (*p*=0.22, *d*=-0.07 [-0.19, 0.04], *BF*_*10*_=0.14). None of the conclusions materially changed when mean-centered study, age, and biological sex were included as nuisance variates. In sum, at least when viewed through the lens of hemodynamics and the MTC paradigm, the BST and Ce show statistically indistinguishable responses to certain- and uncertain-threat anticipation.

**Figure 4.**
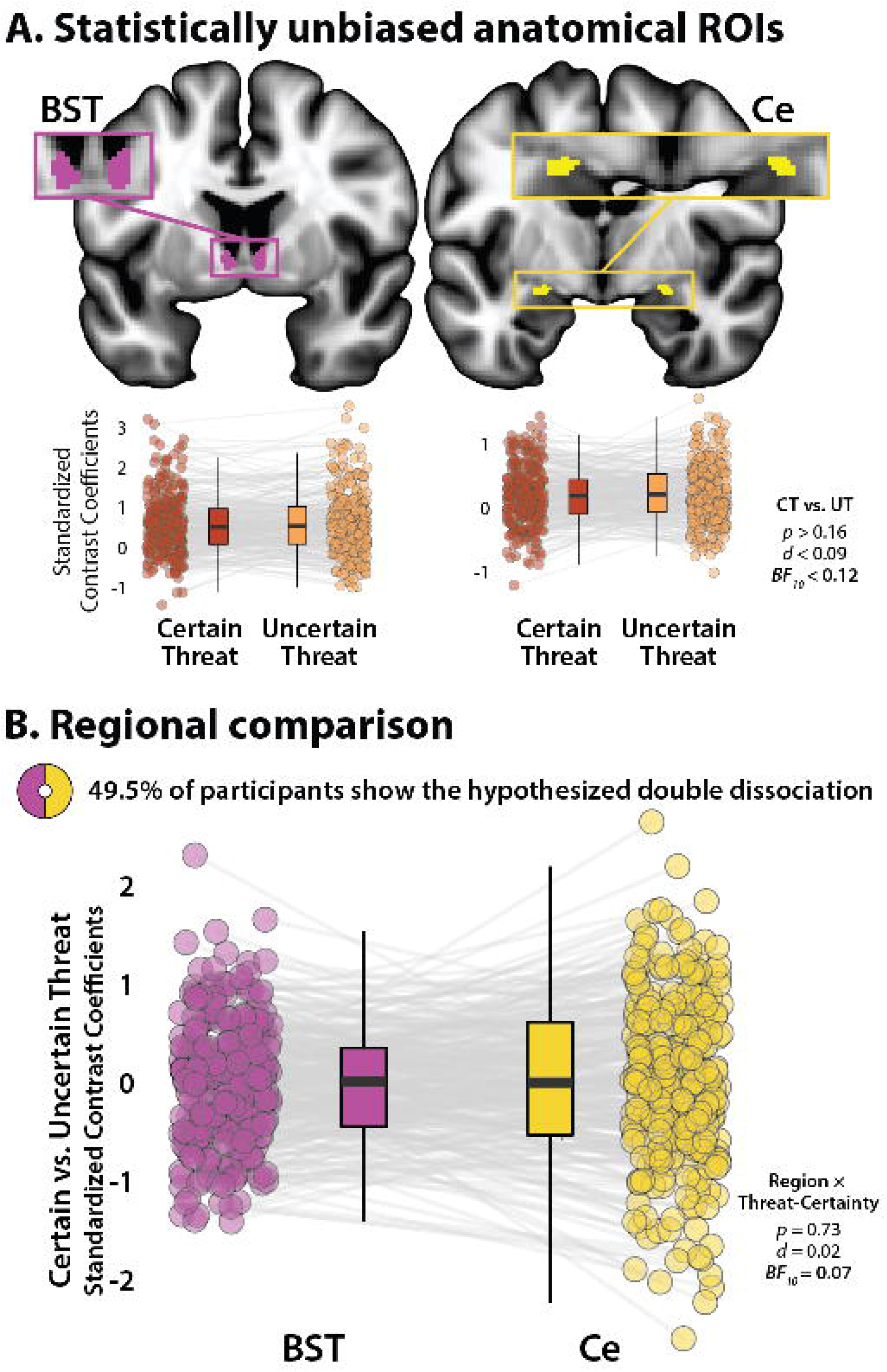
The human BST and Ce show statistically indistinguishable responses during certain- and uncertain-threat anticipation. **(A) *Anatomical ROIs*.** Probabilistic anatomical ROIs provided statistically unbiased estimates of BST and Ce activation during certain- and uncertain-threat anticipation. Leveraging spatially unsmoothed data, regression coefficients were extracted and averaged across voxels for each combination of ROI, task contrast, and participant. Box plots underscore negligible activation differences during the anticipation of certain-versus-uncertain threat in both the BST (*left*) and the Ce (*right*), contrary to double-dissociation models (*p*>0.16, *d*<0.09, *BF*_*10*_<0.12). Note: The *y*-axis scale differs across ROIs. **(B) *Regional comparison***. A standard repeated-measures GLM was used to directly assess potential regional differences in reactivity to certain-versus-uncertain threat. Contrary to the double-dissociation model, the Region × Threat-Certainty interaction was not significant. Boxplot depicts the interaction as a 1-*df* contrast, that is, the ‘difference of differences.’ Participants were just as likely as not (49.5% vs. 50.5%) to show the hypothesized double dissociation, *Z*_*Sign*_=0.12, *p*=0.91. Boxplots depict the median (*horizontal lines*), interquartile range (*boxes*), and individual participants (*dots*) for each contrast. Whiskers indicate 1.5× the interquartile range. Gray lines depict the sign and magnitude of intra-individual mean differences. Inset ring plot depicts the percentage of participants showing the hypothesized dissociation of regional reactivity to threat (BST: Certain < Uncertain Threat; Ce: Certain > Uncertain Threat). Abbreviations—BF, Bayes Factor; BST, bed nucleus of the stria terminalis; Ce, central nucleus of the amygdala; CT, certain-threat anticipation; *d*, Cohen’s *d*. fMRI, functional magnetic resonance imaging; FWE, family-wise error; GLM, general linear model; ROI, region-of-interest; UT, uncertain-threat anticipation.

## DISCUSSION

In the realm of human neuroimaging research, the present results provide some of the strongest evidence to date that the functional architecture of the EA does not conform to popular double-dissociation models (**Supplementary Figure S1 and Note 1**; Avery et al., 2016; Klumpers et al., 2017; LeDoux & Pine, 2016; NIMH, 2011; Somerville et al., 2013). The Ce and BST are both engaged during periods of threat anticipation, and the degree of engagement is independent of the temporal certainty of threat encounters. In a head-to-head comparison, the Ce and BST showed statistically indistinguishable selectivity for the two kinds of threat (*d*=0.02), with strong Bayesian evidence against regional differences (*BF*_*10*_=0.07; *H*_*0*_ is 14.3 times more likely than *H*_*1*_).

It is worth considering two potential methodological explanations for this null observation. First, might it reflect inadequate EA signal quality (e.g., low signal-to-noise ratio [SNR])? This is unlikely. Our results indicate that we have sufficient statistical power and SNR to detect threat-related activation in the Ce and BST and to detect other types of regional activation differences (i.e., greater BST sensitivity to anticipated threat). Second, might it reflect an artifact of our hemodynamic model? After all, our conclusions are based on a simplified ‘boxcar’ model that assumes static, time-invariant neural responses to anticipated threats. This approach precludes inferences about regional differences in more fleeting activation dynamics, and leaves open the possibility that the Ce and BST differ in their phasic responses. Yet this, too, is unlikely. Leveraging a subset of the present sample (*n*=220) and more sophisticated hemodynamic models, Cornwell and colleagues found statistically indistinguishable responses to anticipated threat, manifesting as sustained levels of heightened Ce/BST activation to temporally uncertain-and-distal encounters and phasic surges of Ce/BST activation to certain-and-imminent encounters (Cornwell et al., 2025). Taken together, this pattern of results reinforces the conclusion that the Ce and BST respond similarly to temporally certain and uncertain threat.

The similarities in Ce and BST function identified here are consistent with other evidence. Both regions are characterized by similar anatomical connectivity, cellular composition, neurochemistry, and gene expression (Fox, Oler, Tromp, et al., 2015). Both are poised to trigger behavioral, psychophysiological, and neuroendocrine responses to threat via dense projections to downstream effector regions (Fox, Oler, Tromp, et al., 2015). Activity in both regions has been shown to co-vary with individual differences in trait anxiety in large-scale studies of nonhuman primates (*n*=592) (Fox, Oler, Shackman, et al., 2015). Among humans, the Ce and BST are recruited by a broad spectrum of threatening and aversive stimuli (Fox & Shackman, 2019; Hur et al., 2020; Shackman & Fox, 2021) and both regions show hyper-reactivity to emotional tasks in individuals with anxiety disorders (Chavanne & Robinson, 2021; Shackman & Fox, 2021). Mechanistic work in rodents demonstrates that microcircuits within and between the Ce and BST are critical governors of defensive responses to both certain and uncertain threats (Chen et al., 2022; Fox & Shackman, 2019; Holley & Fox, 2022; Lange et al., 2017; Moscarello & Penzo, 2022; Pomrenze, Giovanetti, et al., 2019; Pomrenze, Tovar-Diaz, et al., 2019; Ren et al., 2022; Ressler et al., 2020; Zhu et al., 2024). In fact, work in mice using a variant of the MTC shows that projections from the Ce to the BST are necessary for mounting defensive responses to temporally uncertain threat (Lange et al., 2017), dovetailing with the present results. Although our understanding remains far from complete, this body of work underscores the need to reformulate conceptual models that posit a strict functional segregation of certain and uncertain threat processing in the EA.

These observations do not mean that the Ce and BST are functionally identical or interchangeable. Indeed, our results indicate that the BST is more strongly recruited by both kinds of anticipated threat. Work in monkeys demonstrates that BST activity is more closely related to heritable variation (‘nature’) in trait anxiety, whereas Ce activity is more closely related to the variation in trait anxiety that is explained by differences in early-life experience (‘nurture’) (Fox, Oler, Shackman, et al., 2015). The BST also appears to be more closely involved in organizing persistent signs of fear and anxiety following threat encounters (Duvarci et al., 2009; Shackman et al., 2017). Among humans, individual differences in neuroticism/negative emotionality, a prominent risk factor for anxiety disorders and depression, is selectively associated with heightened BST reactivity to uncertain-threat anticipation, an association that remains evident when controlling for Ce reactivity (Grogans et al., 2024). Clarifying the nature of these regional differences is an important avenue for future research. This endeavor is likely to benefit from the use of pharmacological challenges (e.g., acute benzodiazepine; Grillon & Ernst, 2020; Miles et al., 2011) and theory-driven computational modeling (Holley & Fox, 2022), approaches that would facilitate the development of coordinated cross-species models of fear and anxiety. When combined with parametric manipulations of specific facets of threat (e.g., duration, probability), computational modeling has the potential to address fundamental questions about the function of threat-sensitive brain regions, clarify inconsistencies across paradigms, and foster a common mathematical framework (*‘lingua franca’*) for integrating research across investigators, readouts, and species (Drzewiecki & Fox, 2024).

The processing of uncertain and certain anticipated threats is not confined to the EA. Whole-brain voxelwise analyses revealed a distributed network that encompasses both frontocortical (MCC, AI/FrO, and dlPFC/FP) and other subcortical regions (PAG) regions (**Figure 3**). And while this network is sensitive to both kinds of threat, with co-localization evident in every key region, direct comparison of the two threat conditions revealed greater frontocortical activation during uncertain-threat anticipation. We previously hypothesized that this could reflect differences in either cognitive load or the intensity of distress and arousal across conditions (Hur et al., 2020). On certain-threat trials, the descending integer stream (‘countdown’) provides an overt index of momentary changes in threat imminence. On uncertain-threat trials, this cognitive scaffolding is absent, encouraging reliance on the kinds of sustained, endogenous representations that are the hallmark of frontocortical regions (Tardiff & Curtis, 2025). A second notable difference between the two threat conditions is the greater intensity of distress and arousal elicited by uncertain threat (**Figure 2**), potentially reflecting differences in the cumulative hazard rate (i.e., the mathematical probability of encountering threat, given that it has not yet occurred) across the two threat conditions (Holley et al., 2024). From this perspective, increased frontocortical activation could reflect either heightened fear/anxiety or stronger recruitment of compensatory processes aimed at downregulating negative affect. On the one hand, there is ample evidence that frontocortical regions, including the MCC and AI/FrO, are recruited by a wide variety of aversive challenges, consistent with a role in *generating* negative affect (Bo et al., 2024; Chavanne & Robinson, 2021; Radua et al., 2025; Xu et al., 2020). Of course, they also play a role in *regulating* distress (Bo et al., 2024). In laboratory studies of prompted cognitive reappraisal, MCC and dlPFC/FP activation scales with the degree of regulatory success (Bo et al., 2024; Urry et al., 2009), raising the possibility that frontocortical activation during aversive laboratory challenges partially reflects spontaneous efforts to downregulate or inhibit distress (‘implicit’ regulation; Shackman & Lapate, 2018). Consistent with this hypothesis, we recently showed that heightened MCC and FrO reactivity to the MTC paradigm is associated with dampened emotional reactivity to real-world stressors, indexed using ecological momentary assessment (Hur et al., 2022), an observation that is consistent with evidence that loss of MCC function is associated with increased (‘dysregulated’) emotional reactivity to painful stimuli in humans and amplified defensive responses to threat in monkeys (Davis et al., 1994; Greenspan et al., 2008; Rahman et al., 2021).

Clearly, several challenges remain for the future. First, it will be important to determine whether our conclusions generalize to more demographically representative samples, other types of experimental threat (e.g., social), other kinds of uncertainty (e.g., probability, risk, ambiguity), and more naturalistic paradigms that span longer and more ecologically valid periods of threat anticipation (Mobbs et al., 2021; Wang et al., 2005). It merits comment that the absence of reward trials precludes strong inferences about valence. Second, the Ce and BST are complex and can be subdivided into multiple subdivisions, each containing intermingled cell types with distinct, even opposing functional roles (e.g., anxiogenic vs. anxiolytic) (Fox & Shackman, 2019, 2024; Holley & Fox, 2022; Moscarello & Penzo, 2022). Animal models will be critical for generating testable hypotheses about the most relevant molecules, cell types, and microcircuits (Fox & Shackman, 2019, 2024; Kamboj et al., 2024). Third, fear and anxiety reflect the coordinated interactions of widely distributed neural networks (Liu et al., 2024; Tovote et al., 2015). Moving forward, it will be important to clarify the relevance of functional connectivity within and beyond the EA.

Anxiety disorders impose a staggering burden on global health, afflicting ~360 million individuals annually (GBD2021, 2024). Existing treatments were developed decades ago and have limited efficacy, acceptability, durability, and tolerability (Cuijpers et al., 2024; De Crescenzo et al., 2024; Singewald et al., 2023). Rising to this challenge requires the development of more accurate models of the neural systems governing fear and anxiety in health and disease. Leveraging a well-powered mega-analytic sample, translationally relevant fMRI paradigm, and best-practices analytic approach, the present results demonstrate that the EA systems recruited by certain and uncertain threat are not categorically different, with clear evidence of functional colocalization—*not* segregation—in the Ce and BST. These observations provide an empirically grounded framework for conceptualizing fear and anxiety, for understanding the functional neuroanatomy of threat processing in humans, and for accelerating the development of improved biological interventions for the suffering caused by extreme fear and anxiety.

## ACKNOWLEDGMENTS

Authors acknowledge assistance and critical feedback from D. Bradford, J. Curtin, L. Friedman, C. Lejuez, B. Nacewicz, L. Pessoa, S. Rose, members of the Affective and Translational Neuroscience laboratory, the staff of the Maryland Neuroimaging Center, and the Office of the Registrar at the University of Maryland.

## REFERENCES

Ahrens, S., Wu, M. V., Furlan, A., Hwang, G. R., Paik, R., Li, H., Li, B. (2018). A central extended amygdala circuit that modulates anxiety. J Neurosci, 38, 5567–5583. 10.1523/JNEUROSCI.0705-18.2018

APA. (2022). Diagnostic and statistical manual of mental disorders, text revision (DSM-5-TR) (5 ed.). American Psychiatric Publishing.

Avery, S. N., Clauss, J. A., & Blackford, J. U. (2016). The human BNST: Functional role in anxiety and addiction. Neuropsychopharmacology, 41, 126–141. 10.1038/npp.2015.185

Blanchard, D. C., & Canteras, N. S. (2024). In search of the behavioral and neural basis for differentiating fear and anxiety. Biological Psychiatry: Global Open Science, 4, 394–395. 10.1016/j.bpsgos.2023.05.008

Bo, K., Kraynak, T. E., Kwon, M., Sun, M., Gianaros, P. J., & Wager, T. D. (2024). A systems identification approach using Bayes factors to deconstruct the brain bases of emotion regulation. Nat Neurosci, 27, 975–987. 10.1038/s41593-024-01605-7

Bruzsik, B., Biro, L., Zelena, D., Sipos, E., Szebik, H., Sarosdi, K. R., Toth, M. (2021). Somatostatin neurons of the bed nucleus of stria terminalis enhance associative fear memory consolidation in mice. J Neurosci, 41, 1982–1995. 10.1523/jneurosci.1944-20.2020

Chavanne, A. V., & Robinson, O. J. (2021). The overlapping neurobiology of adaptive and pathological anxiety: a meta-analysis of functional neural activation. American Journal of Psychiatry, 178, 156-164. 10.1176/appi.ajp.2020.19111153

Chen, W. H., Lien, C. C., & Chen, C. C. (2022). Neuronal basis for pain-like and anxiety-like behaviors in the central nucleus of the amygdala. Pain, 163(3), e463–e475. 10.1097/j.pain.0000000000002389

Choi, J. M., Padmala, S., & Pessoa, L. (2012). Impact of state anxiety on the interaction between threat monitoring and cognition. Neuroimage, 59, 1912–1923.

Cohen, J. (1988). Statistical power analysis for the behavioral sciences (2nd ed.). Lawrence Erlbaum Associates.

Cohen, J. R. (1994). The earth is round (p < .05). American Psychologist, 49, 997–1003. 10.1037/0003-066X.49.12.997

Cornwell, B. R., Didier, P. R., Grogans, S. E., Anderson, A. S., Islam, S., Kim, H. C., Shackman, A. J. (2025). A shared threat-anticipation circuit is dynamically engaged at different moments by certain and uncertain threat. Journal of Neuroscience, 45, e2113242025. 10.1523/JNEUROSCI.2113-24.2025

Costafreda, S. G. (2009). Pooling FMRI data: meta-analysis, mega-analysis and multi-center studies. Front Neuroinform, 3, 33. 10.3389/neuro.11.033.2009

Cuijpers, P., Miguel, C., Ciharova, M., Harrer, M., Basic, D., Cristea, I. A., Karyotaki, E. (2024). Absolute and relative outcomes of psychotherapies for eight mental disorders: a systematic review and meta-analysis. World Psychiatry, 23, 267–275. 10.1002/wps.21203

Daniel-Watanabe, L., & Fletcher, P. C. (2022). Are fear and anxiety truly distinct? Biological Psychiatry Global Open Science, 2, 341–349. 10.1016/j.bpsgos.2021.09.006

Davis, K. D., Hutchison, W. D., Lozano, A. M., & Dostrovsky, J. O. (1994). Altered pain and temperature perception following cingulotomy and capsulotomy in a patient with schizoaffective disorder. Pain, 59, 189–199. 0304-3959(94)90071-X [pii]

De Crescenzo, F., De Giorgi, R., Garriga, C., Liu, Q., Fazel, S., Efthimiou, O., Cipriani, A. (2024). Real-world effects of antidepressants for depressive disorder in primary care: population-based cohort study. Br J Psychiatry, 226, 1–10. 10.1192/bjp.2024.194

Drzewiecki, C. M., & Fox, A. S. (2024). Understanding the heterogeneity of anxiety using a translational neuroscience approach. Cogn Affect Behav Neurosci, 24, 228–245. 10.3758/s13415-024-01162-3

Duvarci, S., Bauer, E. P., & Paré, D. (2009). The bed nucleus of the stria terminalis mediates inter-individual variations in anxiety and fear. J Neurosci, 29, 10357–10361. https://doi.org/29/33/10357 [pii] 10.1523/JNEUROSCI.2119-09.2009

Faul, F., Erdfelder, E., Lang, A.-G., & Buchner, A. (2007). G*Power 3: A flexible statistical power analysis program for the social, behavioral, and biomedical sciences. Behavior Research Methods, 39, 175–191.

Fox, A. S., Lapate, R. C., Davidson, R. J., & Shackman, A. J. (2018). The nature of emotion: A research agenda for the 21st century. In A. S. Fox, R. C. Lapate, A. J. Shackman, & R. J. Davidson (Eds.), The nature of emotion. Fundamental questions (2nd ed., pp. 403–417). Oxford University Press.

Fox, A. S., Oler, J. A., Shackman, A. J., Shelton, S. E., Raveendran, M., McKay, D. R., Kalin, N. H. (2015). Intergenerational neural mediators of early-life anxious temperament. Proceedings of the National Academy of Sciences USA, 112, 9118–9122.

Fox, A. S., Oler, J. A., Tromp, D. P., Fudge, J. L., & Kalin, N. H. (2015). Extending the amygdala in theories of threat processing. Trends Neurosci, 38, 319–329. 10.1016/j.tins.2015.03.002

Fox, A. S., & Shackman, A. J. (2019). The central extended amygdala in fear and anxiety: Closing the gap between mechanistic and neuroimaging research. Neuroscience letters, 693, 58–67. 10.1016/j.neulet.2017.11.056

Fox, A. S., & Shackman, A. J. (2024). An honest reckoning with the amygdala and mental illness. American Journal of Psychiatry, 181, 1059–1075. 10.1176/appi.ajp.20240941

Freud, S., Strachey, J., Freud, A., Rothgeb, C. L., & Richards, A. (1959). The standard edition of the complete psychological works of Sigmund Freud, Volume 20 (1925-1926): An autobiographical study, inhibitions, symptoms and anxiety, the question of lay analysis and other works. Hogarth Press.

GBD2021. (2024). Global incidence, prevalence, years lived with disability (YLDs), disability-adjusted life-years (DALYs), and healthy life expectancy (HALE) for 371 diseases and injuries in 204 countries and territories and 811 subnational locations, 1990-2021: a systematic analysis for the Global Burden of Disease Study 2021. The Lancet, 403, 2133–2161. 10.1016/S0140-6736(24)00757-8

Greenspan, J. D., Coghill, R. C., Gilron, I., Sarlani, E., Veldhuijzen, D. S., & Lenz, F. A. (2008). Quantitative somatic sensory testing and functional imaging of the response to painful stimuli before and after cingulotomy for obsessive-compulsive disorder (OCD) [10.1016/j.ejpain.2008.01.007]. European Journal of Pain, 12, 990–999. 10.1016/j.ejpain.2008.01.007

Griessner, J., Pasieka, M., Bohm, V., Grossl, F., Kaczanowska, J., Pliota, P., Haubensak, W. (2021). Central amygdala circuit dynamics underlying the benzodiazepine anxiolytic effect. Mol Psychiatry, 26, 534–544. 10.1038/s41380-018-0310-3

Grillon, C., & Ernst, M. (2020). A way forward for anxiolytic drug development: Testing candidate anxiolytics with anxiety-potentiated startle in healthy humans. Neurosci Biobehav Rev, 119, 348–354. 10.1016/j.neubiorev.2020.09.024

Grogans, S. E., Bliss-Moreau, E., Buss, K. A., Clark, L. A., Fox, A. S., Keltner, D., Shackman, A. J. (2023). The nature and neurobiology of fear and anxiety: State of the science and opportunities for accelerating discovery. Neuroscience & Biobehavioral Reviews, 151, 105237. 10.1016/j.neubiorev.2023.105237

Grogans, S. E., Hur, J., Barstead, M. G., Anderson, A. S., Islam, S., Kuhn, M., Shackman, A. J. (2024). Neuroticism/negative emotionality is associated with increased reactivity to uncertain threat in the bed nucleus of the stria terminalis, not the amygdala. Journal of Neuroscience, 44, e1868232024.

Gronau, Q. F., Ly, A., & Wagenmakers, E.-J. (2020). Informed Bayesian t-Tests. The American Statistician, 74, 137–143. 10.1080/00031305.2018.1562983

Grupe, D. W., & Nitschke, J. B. (2013). Uncertainty and anticipation in anxiety: an integrated neurobiological and psychological perspective. Nat Rev Neurosci, 14, 488–501. https://doi.org/nrn3524 [pii] 10.1038/nrn3524

Grupe, D. W., Wielgosz, J., Davidson, R. J., & Nitschke, J. B. (2016). Neurobiological correlates of distinct post-traumatic stress disorder symptom profiles during threat anticipation in combat veterans. Psychol Med, 46, 1885–1895. 10.1017/S0033291716000374

Gungor, N. Z., & Paré, D. (2016). Functional heterogeneity in the bed nucleus of the stria terminalis. Journal of Neuroscience, 36, 8038–8049.

Holley, D., & Fox, A. S. (2022). The central extended amygdala guides survival-relevant tradeoffs: Implications for understanding common psychiatric disorders. Neuroscience & Biobehavioral Reviews, 142, 104879. 10.1016/j.neubiorev.2022.104879

Holley, D., Varga, E. A., Boorman, E. D., & Fox, A. S. (2024). Temporal dynamics of uncertainty cause anxiety and avoidance. Computational Psychiatry, 8, 85–91. 10.5334/cpsy.105

Hur, J., Kuhn, M., Grogans, S. E., Anderson, A. S., Islam, S., Kim, H. C., Shackman, A. J. (2022). Anxiety-related frontocortical activity is associated with dampened stressor reactivity in the real world. Psychological Science, 33, 906–924. 10.1101/2021.03.17.435791

Hur, J., Smith, J. F., DeYoung, K. A., Anderson, A. S., Kuang, J., Kim, H. C., Shackman, A. J. (2020). Anxiety and the neurobiology of temporally uncertain threat anticipation. Journal of Neuroscience, 40, 7949–7964. 10.1101/2020.02.25.964734

Kamboj, S., Calrson, E. L., Ander, B. P., Hanson, K. L., Murray, K. D., Fudge, J. L., Fox, A. S. (2024). Translational Insights from cell-type variation across amygdala subnuclei in rhesus monkeys and mumans. American Journal of Psychiatry, 181, 1086–1102.

Kim, H. C., Kaplan, C. M., Islam, S., Anderson, A. S., Piper, M. E., Bradford, D. E., Shackman, A. J. (2023). Acute nicotine abstinence amplifies subjective withdrawal symptoms and threat-evoked fear and anxiety, but not extended amygdala reactivity. PLoS One, 18, e0288544. 10.1371/journal.pone.0288544

Klumpers, F., Kroes, M. C. W., Baas, J., & Fernandez, G. (2017). How human amygdala and bed nucleus of the stria terminalis may drive distinct defensive responses. J Neurosci, 37, 9645–9656. 10.1523/JNEUROSCI.3830-16.2017

Lange, M. D., Daldrup, T., Remmers, F., Szkudlarek, H. J., Lesting, J., Guggenhuber, S., Pape, H. C. (2017). Cannabinoid CB1 receptors in distinct circuits of the extended amygdala determine fear responsiveness to unpredictable threat. Mol Psychiatry, 22, 1422–1430. 10.1038/mp.2016.156

LeDoux, J. E., & Pine, D. S. (2016). Using neuroscience to help understand fear and anxiety: A two-system framework. Am J Psychiatry, 173, 1083–1093. 10.1176/appi.ajp.2016.16030353

Lee, S. C., Amir, A., Haufler, D., & Pare, D. (2017). Differential recruitment of competing valence-related amygdala networks during anxiety. Neuron, 96, 81-88.e85. 10.1016/j.neuron.2017.09.002

Liu, X., Jiao, G., Zhou, F., Kendrick, K. M., Yao, D., Gong, Q., Becker, B. (2024). A neural signature for the subjective experience of threat anticipation under uncertainty. Nat Commun, 15, 1544. 10.1038/s41467-024-45433-6

Marcinkiewcz, C. A., Mazzone, C. M., D’Agostino, G., Halladay, L. R., Hardaway, J. A., DiBerto, J. F., Kash, T.L. (2016). Serotonin engages an anxiety and fear-promoting circuit in the extended amygdala. Nature, 537, 97–101. 10.1038/nature19318

Marvar, P. J., Andero, R., Hurlemann, R., Lago, T. R., Zelikowsky, M., & Dabrowska, J. (2021). Limbic neuropeptidergic modulators of emotion and their therapeutic potential for anxiety and post-traumatic stress disorder. J Neurosci, 41, 901–910. 10.1523/jneurosci.1647-20.2020

Miles, L., Davis, M., & Walker, D. (2011). Phasic and sustained fear are pharmacologically dissociable in rats. Neuropsychopharmacology, 36, 1563–1574. 10.1038/npp.2011.29

Mobbs, D., Wise, T., Suthana, N., Guzmán, N., Kriegeskorte, N., & Leibo, J. Z. (2021). Promises and challenges of human computational ethology. Neuron, 109, 2224–2238. 10.1016/j.neuron.2021.05.021

Moscarello, J. M., & Penzo, M. A. (2022). The central nucleus of the amygdala and the construction of defensive modes across the threat-imminence continuum. Nat Neurosci, 25, 999–1008. 10.1038/s41593-022-01130-5

Murty, D., Song, S., Morrow, K., Kim, J., Hu, K., & Pessoa, L. (2022). Distributed and multifaceted effects of threat and safety. J Cogn Neurosci, 34, 495–516. 10.1162/jocn_a_01807

Murty, D. V. P. S., Song, S., Surampudi, S. G., & Pessoa, L. (2023). Threat and reward imminence processing in the human brain. J Neurosci, 43, 2973–2987. 10.1523/jneurosci.1778-22.2023

Nichols, T., Brett, M., Andersson, J., Wager, T., & Poline, J. B. (2005). Valid conjunction inference with the minimum statistic. Neuroimage, 25, 653–660. https://doi.org/S1053-8119(04)00750-5 [pii]10.1016/j.neuroimage.2004.12.005 [doi]

NIMH. (2011). Negative valence systems: Workshop proceedings (March 13, 2011 – March 15, 2011; Rockville, Maryland). Retrieved July 1 from https://www.nimh.nih.gov/research/research-funded-by-nimh/rdoc/negative-valence-systems-workshop-proceedings.shtml

Petre, B., Kragel, P., Atlas, L. Y., Geuter, S., Jepma, M., Koban, L., Wager, T. D. (2022). A multistudy analysis reveals that evoked pain intensity representation is distributed across brain systems. PLoS biology, 20, e3001620. 10.1371/journal.pbio.3001620

Poldrack, R. A., Baker, C. I., Durnez, J., Gorgolewski, K. J., Matthews, P. M., Munafò, M. R., Yarkoni, T. (2017). Scanning the horizon: towards transparent and reproducible neuroimaging research. Nature Reviews Neuroscience, 18, 115–126.

Poldrack, R. A., Baker, C. I., Durnez, J., Gorgolewski, K. J., Matthews, P. M., Munafo, M. R., Yarkoni, T. (2017). Scanning the horizon: towards transparent and reproducible neuroimaging research. Nat Rev Neurosci, 18, 115–126. 10.1038/nrn.2016.167

Pomrenze, M. B., Giovanetti, S. M., Maiya, R., Gordon, A. G., Kreeger, L. J., & Messing, R. O. (2019). Dissecting the roles of GABA and neuropeptides from rat central amygdala CRF neurons in anxiety and fear learning. Cell Rep, 29, 13–21 e14. 10.1016/j.celrep.2019.08.083

Pomrenze, M. B., Tovar-Diaz, J., Blasio, A., Maiya, R., Giovanetti, S. M., Lei, K., Messing, R. O. (2019). A corticotropin releasing factor network in the extended amygdala for anxiety. J Neurosci, 39, 1030–1043. 10.1523/JNEUROSCI.2143-18.2018

Radua, J., Savage, H. S., Vilajosana, E., Jamieson, A., Abler, B., Åhs, F., Fullana, M. A. (2025). Neural correlates of human fear conditioning and sources of variability: An fMRI mega-analysis and normative modelling study of 2,199 individuals. Nature Communications, 16, 7869.

Rahman, S. S., Mulvihill, K., Wood, C. M., Quah, S. K. L., Horst, N. K., Clarke, H. F., Roberts, A. C. (2021). Differential contribution of anterior and posterior midcingulate subregions to distal and proximal threat reactivity in marmosets. Cerebral Cortex, 31, 4765–4780. 10.1093/cercor/bhab121

Ren, J., Lu, C. L., Huang, J., Fan, J., Guo, F., Mo, J. W., Cao, X. (2022). A distinct metabolically defined central nucleus circuit bidirectionally controls anxiety-related behaviors. J Neurosci, 42, 2356–2370. 10.1523/jneurosci.1578-21.2022

Ressler, R. L., Goode, T. D., Evemy, C., & Maren, S. (2020). NMDA receptors in the CeA and BNST differentially regulate fear conditioning to predictable and unpredictable threats. Neurobiology of Learning and Memory, 174, 107281. 10.1016/j.nlm.2020.107281

Salimi-Khorshidi, G., Smith, S. M., Keltner, J. R., Wager, T. D., & Nichols, T. E. (2009). Meta-analysis of neuroimaging data: a comparison of image-based and coordinate-based pooling of studies. Neuroimage, 45, 810–823. https://doi.org/S1053-8119(08)01290-1 [pii] 10.1016/j.neuroimage.2008.12.039

Salomon, J. A., Haagsma, J. A., Davis, A., de Noordhout, C. M., Polinder, S., Havelaar, A. H., Vos, T. (2015). Disability weights for the Global Burden of Disease 2013 study. Lancet Glob Health, 3, e712–723. 10.1016/S2214-109X(15)00069-8

Schimmack, U. (2019). Statistics wars: Don’t change alpha. Change the null-hypothesis! Retrieved December 15 from https://replicationindex.com/category/nil-hypothesis/

Schmalz, X., Biurrun Manresa, J., & Zhang, L. (2023). What is a Bayes factor? Psychological Methods, 28, 705–718. 10.1037/met0000421

Schmitz, A., & Grillon, C. (2012). Assessing fear and anxiety in humans using the threat of predictable and unpredictable aversive events (the NPU-threat test). Nat Protoc, 7, 527–532. https://doi.org/nprot.2012.001 [pii] 10.1038/nprot.2012.001

Schönbrodt, F. D., Wagenmakers, E. J., Zehetleitner, M., & Perugini, M. (2017). Sequential hypothesis testing with Bayes factors: Efficiently testing mean differences. Psychol Methods, 22, 322–339. 10.1037/met0000061

Shackman, A. J., & Fox, A. S. (2016). Contributions of the central extended amygdala to fear and anxiety. Journal of Neuroscience, 36, 8050–8063. 10.1523/JNEUROSCI.0982-16.2016

Shackman, A. J., & Fox, A. S. (2021). Two decades of anxiety neuroimaging research: New insights and a look to the future. American Journal of Psychiatry, 178, 106–109.

Shackman, A. J., Fox, A. S., Oler, J. A., Shelton, S. E., Oakes, T. R., Davidson, R. J., & Kalin, N. H. (2017). Heightened extended amygdala metabolism following threat characterizes the early phenotypic risk to develop anxiety-related psychopathology. Molecular Psychiatry, 22, 724–732.

Shackman, A. J., Grogans, S. E., & Fox, A. S. (2024). Fear, anxiety, and the functional architecture of the human central extended amygdala. Nature Reviews Neuroscience, 25, 587–588. 10.1038/s41583-024-00832-y

Shackman, A. J., & Lapate, R. C. (2018). How are emotions regulated by context and cognition? In A. S. Fox, R. C. Lapate, A. J. Shackman, & R. J. Davidson (Eds.), The nature of emotion. Fundamental questions (2nd ed., pp. 177–179). Oxford University Press.

Singewald, N., Sartori, S. B., Reif, A., & Holmes, A. (2023). Alleviating anxiety and taming trauma: Novel pharmacotherapeutics for anxiety disorders and posttraumatic stress disorder. Neuropharmacology, 226, 109418. 10.1016/j.neuropharm.2023.109418

Somerville, L. H., Wagner, D. D., Wig, G. S., Moran, J. M., Whalen, P. J., & Kelley, W. M. (2013). Interactions between transient and sustained neural signals support the generation and regulation of anxious emotion. Cereb Cortex, 23, 49–60. 10.1093/cercor/bhr373

Tardiff, N., & Curtis, C. E. (2025). Short-term and working memory. In J. Wixted (Ed.), Learning and memory: A comprehensive reference (3rd ed., pp. 112–133). Academic Press. 10.1016/B978-0-443-15754-7.00025-0

Theiss, J. D., Ridgewell, C., McHugo, M., Heckers, S., & Blackford, J. U. (2017). Manual segmentation of the human bed nucleus of the stria terminalis using 3T MRI. Neuroimage, 146, 288–292. 10.1016/j.neuroimage.2016.11.047

Tillman, R. M., Stockbridge, M. D., Nacewicz, B. M., Torrisi, S., Fox, A. S., Smith, J. F., & Shackman, A. J. (2018). Intrinsic functional connectivity of the central extended amygdala. Human Brain Mapping, 39, 1291–1312.

Tovote, P., Fadok, J. P., & Lüthi, A. (2015). Neuronal circuits for fear and anxiety. Nat Rev Neurosci, 16, 317–331. 10.1038/nrn3945

Urry, H. L., van Reekum, C. M., Johnstone, T., & Davidson, R. J. (2009). Individual differences in some (but not all) medial prefrontal regions reflect cognitive demand while regulating unpleasant emotion. Neuroimage, 47, 852–863. https://doi.org/S1053-8119(09)00581-3 [pii] 10.1016/j.neuroimage.2009.05.069

van Doorn, J., van den Bergh, D., Böhm, U., Dablander, F., Derks, K., Draws, T., Wagenmakers, E.-J. (2021). The JASP guidelines for conducting and reporting a Bayesian analysis. Psychonomic Bulletin & Review, 28, 813–826. 10.3758/s13423-020-01798-5

Wagenmakers, E.-J., Love, J., Marsman, M., Jamil, T., Ly, A., Verhagen, J., Morey, R. D. (2018). Bayesian inference for psychology. Part II: Example applications with JASP. Psychonomic Bulletin & Review, 25, 58–76. 10.3758/s13423-017-1323-7

Wang, J., Rao, H., Wetmore, G. S., Furlan, P. M., Korczykowski, M., Dinges, D. F., & Detre, J. A. (2005). Perfusion functional MRI reveals cerebral blood flow pattern under psychological stress. Proc Natl Acad Sci U S A, 102, 17804–17809. https://doi.org/0503082102 [pii] 10.1073/pnas.0503082102

Xu, A., Larsen, B., Baller, E. B., Scott, J. C., Sharma, V., Adebimpe, A., Satterthwaite, T. D. (2020). Convergent neural representations of experimentally-induced acute pain in healthy volunteers: A large-scale fMRI meta-analysis. Neurosci Biobehav Rev, 112, 300–323. 10.1016/j.neubiorev.2020.01.004

Zhu, Y., Xie, S. Z., Peng, A. B., Yu, X. D., Li, C. Y., Fu, J. Y., Li, X. M. (2024). Distinct circuits from the central lateral amygdala to the ventral part of the bed nucleus of stria terminalis regulate different fear memory. Biol Psychiatry, 95, 732–744. 10.1016/j.biopsych.2023.08.022

