## Supplemental File for "Fear, anxiety, and the extended amygdala—*Absence of evidence for strict functional segregation*"

**Supplementary Materials**

Paige R. Didier^1^

Shannon E. Grogans^1^

Claire M. Kaplan^4^

Hyung Cho Kim^2,3^

Samiha Islam^5^

Allegra S. Anderson^6^

Rachael M. Tillman^7^

Manuel Kuhn^8^

Juyoen Hur^9^

Andrew S. Fox^10,11^

Kathryn A. DeYoung^1^

Jason F. Smith^1^

Alexander J. Shackman^1,2,3^

Department of ^1^Psychology, ^2^Neuroscience and Cognitive Science Program, and ^3^Maryland Neuroimaging Center, University of Maryland, College Park, MD 20742 USA. ^4^CommonSpirit Neuropsychology, Lakewood, CO 80228 USA. ^5^Rees-Jones Center for Foster Care Excellence, Children’s Heath, Dallas, TX 75207 USA. ^6^Department of Psychiatry and Human Behavior, Brown University, Providence, RI 02912 USA. ^7^McGill Neuropsychology, Bethesda, MD 20814 USA. ^8^Institute of Medical Psychology, Charité Universitätsmedizin Berlin, 10117 Berlin, Germany. ^9^Department of Psychology, Yonsei University, Seoul 03722, Republic of Korea. ^10^Department of Psychology and ^11^California National Primate Research Center, University of California, Davis, CA 95616 USA

**Please address manuscript correspondence to**

Alexander J. Shackman

**SUPPLEMENTARY NOTE 1. *Fear, anxiety, and the extended amygdala.***

In this section, we briefly highlight some prominent neuroscientific definitions of fear and anxiety and recent theoretical claims about the functional organization of the two major subdivisions of the extended amygdala (EA), including the central nucleus of the amygdala (Ce/CeA) and the bed nucleus of the stria terminalis (BST/BNST).

By design, this brief survey is meant to be illustrative, *not* comprehensive. For a panel discussion of related conceptual issues, see Grogans and colleagues (Grogans et al., 2023). For an overview of popular assays for assessing fear and anxiety in different species, see Shackman and colleagues (Shackman et al., 2016). For an overview of concerns centered on the imprecision and inconsistency of fear-and-anxiety terminology see Perusini and Fanselow (Perusini & Fanselow, 2015) and Shackman and Fox (Shackman & Fox, 2016).

Avery et al. *Neuropsychopharmacology* 2016 (Avery et al., 2016)

- *“The amygdala mediates short-term, phasic responses to immediate threats, whereas the BNST mediates sustained responses to contextual, diffuse, and unpredictable threats.”*

Daniel-Watanabe & Fletcher *Biological Psychiatry Global Open Science* 2022 (Daniel-Watanabe & Fletcher, 2022)

- *“Fear is related to the presence, or imminent presence, of the aversive stimulus, while anxiety is considered the more protracted state produced by a sustained expectation that the aversive event is likely to occur.”*
- *“Evidence that both amygdala and BNST are responsive to both predictable and unpredictable threats**… calls into question this overall claim for a neural distinction between fear and anxiety.”*
- *“It is difficult to escape the conclusion that the current distinction between fear and anxiety is an unreliable one. While it has been useful in guiding research and clinical work, the inconsistencies suggest that there is a need to reexamine the distinction and consider the importance of other aspects of the experience of anxiety, such as uncertainty and avoidance.”*

Davis et al. *Neuropsychopharmacology* 2010 (Davis et al., 2010)

- *“Although the symptoms of fear and anxiety are very similar, they differ in terms of certain key dimensions….Fear is prompted by imminent and real danger, and galvanizes active defensive responses. In contrast, anxiety is often elicited by less specific and less predictable threats, or by those that are physically or psychologically more distant.”*
- *“The BLA [basolateral amygdala] sends heavy projections to both the CeA and the BNST… The heaviest projections…are to the CeA_M_ [medial division of the Ce/CeA], which in turn projects to the hypothalamus and brainstem to mediate phasic fear responses…A long duration fear stimulus activates the BLA, which then rapidly activates the CeA_M_ to produce phasic fear through the release of glutamate acting on AMPA/kainate receptors on CeA_M_ neurons. Shortly thereafter, inputs to the CeA_L_ [lateral division of the Ce/CeA] then release CRF into the BNST to cause a long-lasting sustained fear reaction. Inhibitory feedback to the CeA_M_ from either the CeA_L_ or the BNST shuts down the CeA_M_, allowing a seamless transition from phasic to sustained fear.*

Domschke *Biological Psychiatry Global Open Science* 2022 (Domschke, 2022)

- *“Fear and anxiety seem to represent closely interrelated diagnostic constructs remaining to be further interrogated for their shared and unique molecular, neuronal, physiological, and behavioral substrates—quite in accordance with Martin Heidegger’s reconciling notion:* “Of course it still remains obscure how [anxiety] is connected ontologically with fear. Obviously these are kindred phenomena.”

Fox & Shackman *Neuroscience Letters* 2019 (Fox & Shackman, 2019)

- *“On balance, the brain imaging literature suggests that the Ce and BST, while certainly not interchangeable, are more alike than different.”*
- *“The central extended amygdala plays a crucial role in evaluating and responding to a broad spectrum of threat-related cues and contexts. While they are certainly not interchangeable, the Ce and the BST show similar patterns of connectivity, cellular composition, neurochemistry, and gene expression. Both are sensitive to uncertain or temporally remote threat; both co-vary with threat-elicited changes in behavior, physiology, and experience; both show phasic responses to acute threat; and both show heightened activity during sustained exposure to diffusely threatening contexts. Work in rodents indicates that both regions play a critical role in organizing sustained defensive responses to a range of potentially threatening cues and contexts.”*

Grupe & Nitschke *Nature Reviews Neuroscience* 2013 (Grupe & Nitschke, 2013)

- *“Fear and anxiety can be distinguished according to how much certainty one has regarding the likelihood, timing or nature of a future threat…Environmental cues indicating the unambiguous presence of an immediate threat give rise to intense 'fearful' defensive behaviours (that is, 'fight or flight'), whereas more diffuse, distal or unpredictable threat cues produce 'anxious' risk assessment behaviour…that is likely to persist until such uncertainty is resolved.”*
- *“We define anxiety…as anticipatory affective, cognitive and behavioural changes in response to uncertainty about a potential future threat.”*

Gungor & Paré *Journal of Neuroscience* 2016 (Gungor & Paré, 2016)

- *Early work stressed the differing involvement of the central amygdala (CeA) and bed nucleus of the stria terminalis (BNST) in the genesis of fear versus anxiety, respectively…This model became extremely influential and now guides a new wave of studies on the role of BNST in humans. Here, we consider evidence for and against this model…This analysis leads us to conclude that BNST's influence is not limited to the generation of anxiety-like responses to diffuse threats, but that it also shapes the impact of discrete threatening stimuli.”*

LeBow & Chen *Molecular Psychiatry* 2016 (Lebow & Chen, 2016)

- *“The amygdala is responsible for mediating specific cue-based fear responses, and thus controls the assessment of immediate or phasic fear.”*
- *“Fear and anxiety can be divided into an immediate threat, for example, the presence of a predator, and pre- or post-encounter threats, for example, the apprehension of encountering a predator again. The BNST is hypothesized to mediate these longer-term responses to anxiety, where a challenge or recovery from a challenge to homeostasis occurs. In contrast, the CeA is thought to mediate shorter duration phasic fear responses, which occur in response to a current challenge to homeostasis.”*

LeDoux & Pine *American Journal of Psychiatry* 2016 (LeDoux & Pine, 2016)

- *“We propose…that the mental state term* fear *be used to describe feelings that occur when the source of harm, the threat, is either immediate or imminent, and* anxiety *be used to describe feelings that occur when the source of harm is uncertain or is distal in space or time.”*
- *“Just as findings demonstrating that the amygdala detects and controls behavioral and physiological responses to immediate threats have supported views of the amygdala as fear-circuit hub, other findings, about responses to uncertain threats, have led to a view of the circuitry of anxiety. Thus, in recent years, animal research has suggested the bed nucleus of the stria terminalis (BNST) is engaged when threats are uncertain…resulting in behavioral inhibition and risk assessment …The BNST has thus come to be for anxiety what the amygdala is for fear—a circuit hub out of which anxious feelings emerge.”*

Mobbs et al. *Trends in Cognitive Sciences* 2020 (Mobbs et al., 2020)

- *“Anxiety: a future-oriented emotional state associated with potential and uncertain threats.”*
- *“Fear: an emotion that is associated with a present and identifiable threat.”*

Moscarello & Penzo *Nature Neuroscience* 2022 (Moscarello & Penzo, 2022)

- *In nature, animals display defensive behaviors that reflect the spatiotemporal distance of threats. Laboratory-based paradigms that elicit specific defensive responses in rodents have provided valuable insight into the brain mechanisms that mediate the construction of defensive modes with varying degrees of threat imminence….High- and low-imminence defensive modes…are mediated at the neural-circuit level within the CeA and its downstream targets.*

NIMH *Negative Valence Systems: Workshop Proceedings* 2011 (NIMH, 2011)

- *“Responses to acute threat (Fear): Activation of the brain’s defensive motivational system to promote behaviors that protect the organism from perceived danger. Normal fear involves a pattern of adaptive responses to conditioned or unconditioned threat stimuli (exteroceptive or interoceptive). Fear can involve internal representations and cognitive processing, and can be modulated by a variety of factors.”*

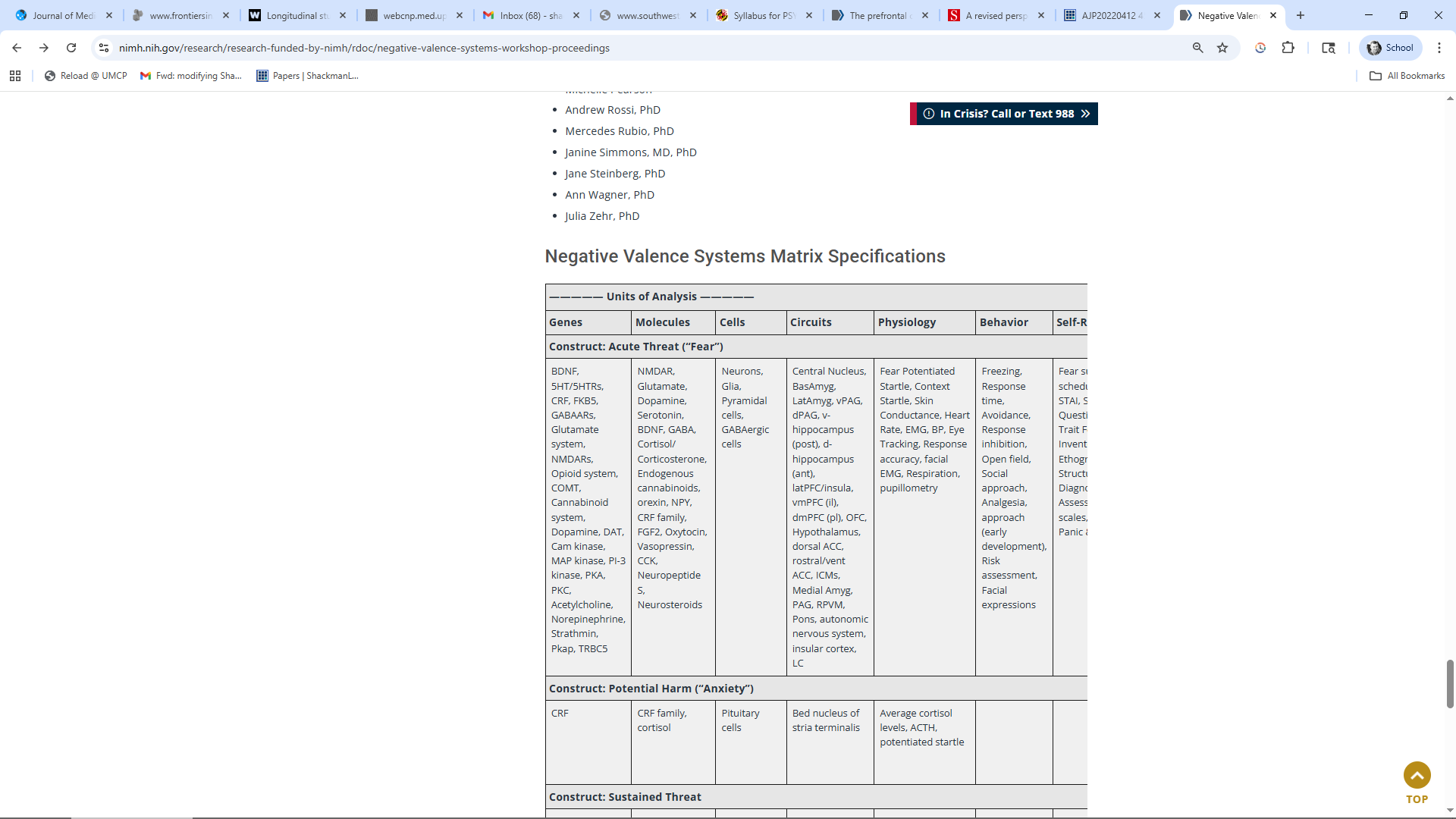

- *“Responses to potential harm (Anxiety): Activation of a brain system in which harm may potentially occur but is distant, ambiguous, or low/uncertain in probability, characterized by a pattern of responses such as enhanced risk assessment (vigilance). These responses to low imminence threats are qualitatively different than the high imminence threat behaviors that characterize fear.”*
- *“While there is considerable overlap in the efferent activity generated, the distinction between these two constructs has found support in both animal and human literatures. In animal models, acute threat is associated with efferent activity from the amygdala, whereas potential harm has been shown to be related to activation of the bed nucleus of the stria terminalis (BNST).”*

Oredu, Lennon, Vervliet & Schiller (Orederu et al., 2024)

- *“Ethological, clinical, and neurobiological evidence strongly suggests that what we colloquially call “fear” is best understood as referring to two different scientific constructs, namely fear and anxiety…Fear is a transient reaction to an identifiable and often proximal threat…and serves to motivate defensive behaviors aimed at coping with the upcoming threat (e.g., fight, flight, freeze). Anxiety, on the other hand, refers to more general, chronic apprehension and worry that is free-floating and not bound to a specific object…Essentially, fear constitutes a state in which a person is actively responding to present danger, while anxiety refers to a person’s response to an uncertain threat that may occur in the future.”*
- *“Inactivation of…BNST selectively impairs innate, but not conditioned, fear responses…These roles are reversed when it comes to conditioned fear, with central nucleus inactivation leading to selective reduction in conditioned fear, and BNST…inactivation sparing conditioned fear…Such divergent findings point to a specific role for the central nucleus in conditioned fear.”*

Tovote et al. *Nature Reviews Neuroscience* 2015 (Tovote et al., 2015)

- *“Whereas fear is evoked by discrete and acutely threaten­ing stimuli, anxiety can be operationalized as an emo­tional response to vague, potential threats.”*

**SUPPLEMENTARY FIGURES AND CAPTIONS**

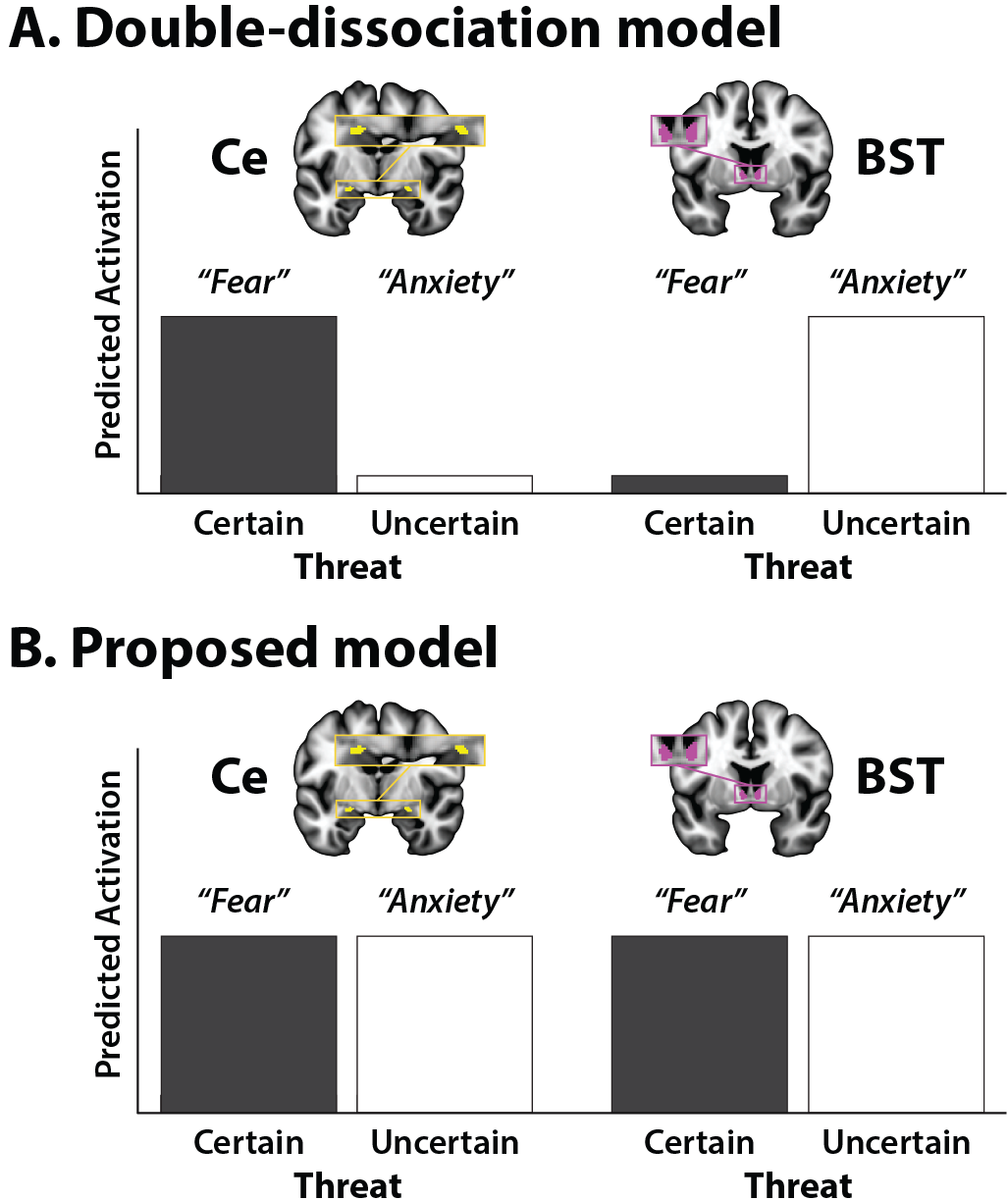

**Supplementary Figure S1.** Conceptual overview of competing hypotheses about the functional architecture of the extended amygdala (EA). **(A)** ***Double-dissociation model.*** Several influential models suggest that fear and anxiety arise from two strictly segregated, doubly dissociable neural systems, positing that the Ce is sensitive to certain (but not uncertain) threat and promotes signs and symptoms of fear; whereas the BST is sensitive to uncertain (but not certain) threat and promotes anxiety (see **Supplementary Note 1**, above). **(B)** ***Proposed model.*** We propose a competing hypothesis, positing that that the Ce and BST play a role in governing responses to both kinds of threat (Fox & Shackman, 2019; Shackman & Fox, 2016). Note: This figure is purely for heuristic purposes and does not depict actual data. Abbreviations—BST, bed nucleus of the stria terminalis; Ce, central nucleus of the amygdala.

**SUPPLEMENTARY METHOD**

**Threat-Anticipation Paradigm**

***Paradigm Structure and Design Considerations.*** The Maryland Threat Countdown paradigm is a well-established, fMRI-optimized variant of temporally uncertain-threat assays that have been validated in rodents and humans (Daldrup et al., 2015; Hefner et al., 2013; Lange et al., 2017; Miles et al., 2011; Moberg et al., 2017).

As shown schematically in **Figure 1**, the paradigm takes the form of a 2 (*Valence:* Threat/Safety) × 2 (*Temporal Certainty:* Uncertain/Certain) randomized, event-related, repeated-measures design (3 scans; 6 trials/condition/scan). Participants were completely informed about the task design and contingencies prior to scanning. Simulations were used to optimize the detection and deconvolution of task-related hemodynamic signals. Stimulus presentation was controlled using Presentation software (version 19.0, Neurobehavioral Systems, Berkeley, CA).

On Certain-Threat trials, participants saw a descending stream of integers (‘count-down;’ e.g., 30, 29, 28...3, 2, 1) for 18.75 s. To ensure robust distress and arousal, the anticipation epoch culminated with the presentation of a noxious electric shock, unpleasant photograph (e.g., mutilated body), and thematically related audio clip (e.g., gunshot). Uncertain-Threat trials were similar, but the integer stream was randomized and presented for an uncertain and variable duration (8.75-30.00 s; *M*=18.75 s). Here, participants knew that something aversive was going to occur but had no way of knowing precisely when. Consistent with methodological recommendations (Shackman & Fox, 2016), the mean duration of the anticipation epoch was identical across conditions, ensuring equal measurement precision. The specific duration was chosen to enhance detection of task-related differences in the blood oxygen level-dependent (BOLD) signal (‘activation’) (Henson, 2007) and to allow sufficient time for sustained responses to become evident. Safety trials were similar but terminated with the delivery of emotionally neutral reinforcers (see below). Valence was continuously signaled during the anticipation epoch (‘countdown’) by the background color of the display. Temporal certainty was signaled by the nature of the integer stream. Certain trials always began with the presentation of the number 30. On Uncertain trials, integers were randomly drawn from a near-uniform distribution ranging from 1 to 45 to reinforce the impression that they could be much shorter or longer than Certain trials and to minimize incidental temporal learning (‘time-keeping’). To concretely demonstrate the variable duration of Uncertain trials, during scanning, the first three Uncertain trials featured short (8.75 s), medium (15.00 s), and long (28.75 s) anticipation epochs. To mitigate potential confusion and eliminate mnemonic demands, a lower-case ‘c’ or ‘u’ was presented at the lower edge of the display throughout the anticipatory epoch. White-noise visual masks (3.2 s) were presented between trials to minimize the persistence of visual reinforcers in iconic memory.

Participants were periodically prompted (following the offset of the white-noise visual mask) to rate the intensity of fear/anxiety experienced a few seconds earlier, during the anticipation period of the prior trial, using a 1 (*minimal*) to 4 (*maximal*) scale and an MRI-compatible response pad (MRA, Washington, PA). Each condition was rated once per scan (16.7% trials). Skin conductance was continuously acquired throughout.

***Procedures.*** Prior to scanning, participants practiced an abbreviated version of the paradigm (without electrical stimulation) until they indicated and staff confirmed understanding. Benign and aversive electrical stimulation levels were individually titrated. *Benign Stimulation.* Participants were asked whether they could “reliably detect” a 20 V stimulus and whether it was “at all unpleasant.” If the participant could not detect the stimulus, the voltage was increased by 4 V and the process repeated. If the participant indicated that the stimulus was unpleasant, the voltage was reduced by 4 V and the process was repeated. The final level chosen served as the benign electrical stimulation during the imaging assessment. *Aversive Stimulation.* Participants received a 100 V stimulus and were asked whether it was “as unpleasant as you are willing to tolerate”—an instruction specifically chosen to maximize anticipatory distress and arousal. If the participant indicated that they were willing to tolerate more intense stimulation, the voltage was increased by 10 V and the process repeated. If the participant indicated that the stimulus was too intense, the voltage was reduced by 5 V and the process repeated. The final level chosen served as the aversive electrical stimulation during the imaging assessment. Following each scan, staff re-assessed whether stimulation was sufficiently intense and increased the level as necessary (for descriptive statistics, see Grogans et al., 2024; Kim et al., 2023).

***Electrical Stimuli.*** Electrical stimuli (100 ms; 2 ms pulses every 10 ms) were generated using an MRI-compatible constant-voltage stimulator system (STMEPM-MRI; Biopac Systems, Inc., Goleta, CA) and delivered using MRI-compatible, disposable carbon electrodes (Biopac) attached to the fourth and fifth digits of the left hand.

***Visual Stimuli.*** Seventy-two aversive and benign photographs (1.8 s) were selected from the International Affective Picture System (for details, see Hur et al., 2020). Visual stimuli were back-projected (Powerlite Pro G5550, Epson America, Inc., Long Beach, CA) onto a semi-opaque screen mounted at the head-end of the scanner bore and viewed using a mirror mounted on the head-coil.

***Auditory Stimuli.*** Seventy-two aversive and benign auditory stimuli (0.8 s) were adapted from open-access online sources and delivered using an amplifier (PA-1 Whirlwind) with in-line noise-reducing filters and ear buds (S14; Sensimetrics, Gloucester, MA) fitted with noise-reducing ear plugs (Hearing Components, Inc., St. Paul, MN).

***Skin Conductance.*** Skin conductance was continuously acquired during each scan using a Biopac system (MP-150; Biopac Systems, Inc., Goleta, CA). Skin conductance (250 Hz; 0.05 Hz high-pass) was measured using MRI-compatible disposable electrodes (EL507) attached to the second and third digits of the left hand.

Data were acquired using a single Siemens Magnetom TIM Trio 3 Tesla scanner (32-channel head-coil). Foam inserts were used to immobilize the participant’s head within the head-coil. Participants were continuously monitored using an eye-tracker (Eyelink 1000; SR Research, Ottawa, Ontario, Canada) and the AFNI real-time motion plugin (Cox, 1996). Eye-tracking data were not recorded. Sagittal T1-weighted anatomical images were acquired using a magnetization prepared rapid acquisition gradient echo sequence (TR=2,400 ms; TE=2.01 ms; inversion time=1,060 ms; flip=8°; slice thickness=0.8 mm; in-plane=0.8×0.8 mm; matrix=300×320; field-of-view=240×256). A T2-weighted image was collected co-planar to the T1-weighted image (TR=3,200 ms; TE=564 ms; flip angle=120°). A multi-band sequence was used to collect oblique-axial echo-planar imaging (EPI) volumes (multiband acceleration=6; TR=1,250 ms; TE=39.4 ms; flip=36.4°; slice thickness=2.2 mm, number of slices=60; in-plane resolution=2.1875×2.1875 mm; matrix=96×96). Data were collected in the oblique-axial plane (approximately −20° relative to the AC-PC plane) to minimize susceptibility artifacts. Three 478-volume EPI scans were acquired. The scanner automatically discarded the first 7 volumes. To enable fieldmap correction, two oblique-axial spin echo (SE) images were collected in opposing phase-encoding directions (rostral-to-caudal and caudal-to-rostral) at the same location and resolution as the functional volumes (i.e., co-planar; TR=7,220 ms; TE=73 ms). Respiration and pulse were continuously acquired during scanning using a respiration belt and photo-plethysmograph affixed to the first digit of the non-dominant hand.

**Skin Conductance Data Processing Pipeline**

Skin conductance data were processed using *PsPM* (version 4.0.2) and in-house Matlab (version 9.9.0.1467703) code (Bach et al., 2018; Bach & Friston, 2013). Data were orthogonalized with respect to pulse and respiration signals and de-spiked using *filloutliers* (150-sample moving-median widow; modified Akima cubic Hermite interpolation). Each scan was then band-pass filtered (0.009-0.333 Hz), median centered, and down-sampled (4 Hz). Participant-specific skin conductance response functions (SCRFs) were estimated by fitting the four parameters of the canonical SCRF (Bach et al., 2010) to the grand-average reinforcer response using *fmincon* and a cost function that maximized variance explained and penalized negative coefficients.

**Skin Conductance Modeling**

Robust general linear models (GLMs) were used to separate electrodermal signals associated with threat anticipation from those evoked by other aspects of the task (e.g., reinforcer presentation). Modeling was performed separately for each participant and scan using *robustfit*. Subject-specific SCRFs were convolved with rectangular regressors time-locked to the presentation of the reinforcers (separately for each trial type), visual masks, and rating prompts. The first-level residuals were then averaged separately for each participant and condition, enabling us to quantify skin conductance level (SCL) during the anticipation (‘countdown’) epochs.

**MRI Pipeline**

Methods were optimized to minimize spatial normalization error and other potential sources of noise and are similar to other recent work by our group (Cornwell et al., 2025). Data were visually inspected before and after processing for quality assurance.

***Anatomical Data Processing.*** T1- and T2-weighted images were inhomogeneity corrected using *N4* (Tustison et al., 2010) and denoised using *ANTS* (Avants et al., 2011). The brain was then extracted using a combination of *BEaST* (Eskildsen et al., 2012) and brain-extracted and normalized reference brains from *IXI* (BIAC, 2022). Extracted T1 images were normalized to a variant of the 1-mm T1-weighted MNI152 template that was modified to remove extracerebral tissue (non-linear 6th-generation symmetric average; Grabner et al., 2006). Normalization was performed using the diffeomorphic approach implemented in *SyN* (version 2.3.4) (Avants et al., 2011). T2-weighted images were rigidly co-registered with the corresponding T1 prior to normalization. The brain extraction mask from the T1 was then applied. Tissue priors were unwarped to native space using the inverse diffeomorphic transformation (Lorio et al., 2016). Brain-extracted T1 and T2 images were segmented using native-space priors generated in *FAST* (version 6.0.4) for use in T1-EPI co-registration (Jenkinson et al., 2012).

***Fieldmap Data Processing.*** SE images and *topup* were used to create fieldmaps. Fieldmaps were converted to radians, median-filtered, and smoothed (2-mm). The average of the distortion-corrected SE images was inhomogeneity corrected using *N4* and masked to remove extracerebral voxels using *3dSkullStrip* (version 19.1.00). The resulting mask was minimally eroded to further exclude extracerebral voxels.

***Functional Data Processing.*** EPI files were de-spiked using *3dDespike*, slice-time corrected to the TR-center using *3dTshift*, and motion-corrected to the first volume and inhomogeneity corrected using *ANTS* (12-parameter affine). Transformations were saved in ITK-compatible format for subsequent processing (McCormick et al., 2014). The first volume was extracted for EPI-T1 co-registration. The reference EPI volume was simultaneously co-registered with the corresponding T1-weighted image in native space and corrected for geometric distortions using boundary-based registration (Jenkinson et al., 2012). This step incorporated the previously created fieldmap, undistorted SE, T1, white matter (WM) image, and masks. The spatial transformations necessary to transform each EPI volume from native space to the reference EPI, from the reference EPI to the T1, and from the T1 to the template were concatenated and applied to the processed EPI data in a single step to minimize incidental spatial blurring. Normalized EPI data were resampled (2 mm^3^) using fifth-order b-splines. Voxelwise analyses employed data that were spatially smoothed (4-mm) using *3DblurInMask*. To minimize signal mixing, smoothing was confined to the gray-matter compartment, defined using a variant of the Harvard-Oxford cortical and subcortical atlases that was expanded to include the bed nucleus of the stria terminalis (BST) and periaqueductal gray (PAG) (Desikan et al., 2006; Edlow et al., 2012; Frazier et al., 2005; Makris et al., 2006; Theiss et al., 2017). Focal analyses of the extended amygdala (EA) leveraged spatially unsmoothed data and anatomically defined regions of interest (ROIs; see below), consistent with prior work by our group (Cornwell et al., 2025).

**fMRI Data Modeling**

***First-Level Modeling.*** For each participant, first-level modeling was performed using GLMs implemented in *SPM12* (version 7771), with the default autoregressive model and the temporal band-pass filter set to the hemodynamic response function (HRF) and 128 s (~0.0078-0.1667 Hz; Wellcome Centre for Human Neuroimaging, 2022). Regressors were convolved with a canonical HRF and its temporal derivative. For the threat-anticipation paradigm, hemodynamic activity was modeled using variable-duration rectangular (‘boxcar’) regressors that spanned the entirety of the anticipation (‘countdown’) epochs of the Uncertain-Threat, Certain-Threat, and Uncertain-Safety trials. To maximize design efficiency, Certain-Safety anticipation served as the reference condition and contributed to the implicit baseline estimate. Epochs corresponding to the presentation of the four types of reinforcers, white-noise visual masks, and rating prompts were simultaneously modeled using the same approach. EPI volumes acquired before the first trial and following the final trial were unmodeled and contributed to the baseline estimate. Consistent with prior work (Cornwell et al., 2025), nuisance variates included volume-to-volume displacement and first derivative, 6 motion parameters and first derivatives, cerebrospinal fluid (CSF) signal, instantaneous pulse and respiration rates, and nuisance signals (e.g., brain edge, CSF edge, global motion, WM, and extracerebral soft tissue) (Anderson et al., 2011; Pruim et al., 2015). Volumes with excessive volume-to-volume displacement (>0.5 mm) and those during and immediately following reinforcer delivery were censored. On average, 3.39 volumes were censored per run (*SD*=4.42).

***Anatomical ROIs.*** Ce and BST activation was quantified using well-established, anatomically defined regions-of-interest (ROIs) and spatially unsmoothed data (Theiss et al., 2017; Tillman et al., 2018). The BST ROI mostly encompasses the supra-commissural BST, given the difficulty of reliably discriminating the sub-commissural BST border in standard anatomical images (Kruger et al., 2015; Walter et al., 1991). Bilateral ROIs were decimated to the 2-mm resolution of the fMRI data. ROI analyses used standardized regression coefficients extracted and averaged for each combination of task contrast (e.g., Uncertain-Threat anticipation vs. Uncertain-Safety anticipation), region, and participant. Anatomical ROIs enable statistically unbiased tests of regional sensitivity to specific experimental manipulations (i.e., Region × Condition effects), including potential single and double dissociations (e.g., BST: Uncertain > Certain Threat; Ce: Uncertain < Certain Threat) (Fox et al., 2018; Poldrack et al., 2017).

**Analytic Strategy**

***Overview.*** Except where noted otherwise, analyses were performed using *SPM12* (version 7771) and *SPSS* (version 27.0.1.0) (Wellcome Centre for Human Neuroimaging, 2022). Diagnostic procedures and data visualizations were used to confirm that test assumptions were satisfied (Tukey, 1977). Standardized frequentist (Cohen’s *d*) effect sizes were interpreted using established benchmarks (Cohen, 1988; Cohen, 1994; Schimmack, 2019), ranging from *large* (*d*=0.80), to *medium* (*d*=0.50), to *small* (*d*=0.20), to *nil* (*d*≤0.10). Some figures were created using created using *ggplot2* (version 3.3.6) (Wickham, 2016) and *MRIcron* (Rorden, 2019). Clusters and local maxima were labeled using the Harvard–Oxford atlas (Desikan et al., 2006; Frazier et al., 2005; Makris et al., 2006), supplemented by other resources (ten Donkelaar et al., 2018).

*JASP* (version 0.16.4.0) was used to compute Bayesian effect sizes for select analyses (Love et al., 2019; van Doorn et al., 2021). Here, Bayes Factor (*BF_10_*) quantifies the relative performance of the null hypothesis (*H_0_*; e.g., the absence of a credible mean difference) and the alternative hypothesis (*H_1_*; e.g., the presence of a credible mean difference), on a 0 to ∞ scale. A key advantage of *BF* is that it can be used to quantify the relative strength of the evidence for *H_0_* (test the null), unlike conventional frequentist null-hypothesis significance tests (Bo et al., 2024; Wagenmakers et al., 2018). It also does not require the data analyst to arbitrarily decide what constitutes a ‘statistically indistinguishable’ difference, in contrast to traditional equivalence tests (Hur et al., 2020). The Bayesian approach provides readily interpretable, principled effect-size benchmarks (van Doorn et al., 2021). Values >1 were interpreted as evidence of mean differences in activation across conditions, ranging from *strong* (*BF_10_*>10), to *moderate* (*BF_10_*=3-10), to *weak* (*BF_10_*=1-3). Values <1 were interpreted as evidence of statistical equivalence (i.e., support for the null hypothesis), ranging from s*trong* (*BF_10_*<0.10), to *moderate* (*BF_10_*=0.10-0.33), to *weak* (*BF_10_*=0.33-1). The reciprocal of *BF_10_* represents the relative likelihood of the null hypothesis (e.g., *BF_10_*=0.10, *H_0_* is 10 times more likely than *H_1_*). Bayesian effects were computed using a noninformative zero-centered Cauchy distribution (*ω*=1/√2), the default setting in *JASP* and the field standard for two-sided tests (Gronau et al., 2020; Schmalz et al., 2023; Schönbrodt et al., 2017; van Doorn et al., 2021; Wagenmakers et al., 2018). Across tests, the estimated error of the MCMC-derived (Markov Chain Monte Carlo) *BF_10_* estimates was negligible (<0.30%) and stable across a range of priors.

***Ratings and Psychophysiology.*** We used repeated-measures general linear models (GLMs) to confirm that the threat-anticipation paradigm amplified subjective symptoms of distress (in-scanner fear/anxiety ratings) and objective signs of arousal (SCL). Interactions were probed using focal contrasts. Sensitivity analyses confirmed that none of the conclusions materially changed when controlling for potential nuisance variation in mean-centered study, age, and assigned sex (for additional details, see the study OSF collection).

***Whole-Brain Voxelwise Tests.*** Spatially smoothed (4-mm) data and whole-brain voxelwise (‘second-level’) repeated-measures GLMs (‘random effects’) were used to compare each threat-anticipation condition to its corresponding control condition (e.g., Uncertain-Threat vs. Uncertain-Safety anticipation), while accounting for potential nuisance variation in mean-centered dummy-coded study (Grogans et al., 2024; Kim et al., 2023), age, and assigned sex. Significance was assessed using *p*<0.05 (whole-brain familywise error [FWE] corrected). A minimum-conjunction test (logical ‘AND’) was used to identify the subset of voxels significantly sensitive to both Certain- *and* Uncertain-Threat anticipation (Nichols et al., 2005). We also directly examined potential differences in anticipatory activity between the two threat conditions (Certain Threat vs. Uncertain Threat). We did not examine hemodynamic responses to reinforcer presentation given the possibility of artifact.

***Anatomical ROIs.*** As a precursor to hypothesis testing, a series of one-sample Student’s *t*-tests was used to confirm that the EA (BST/Ce) ROIs—which leveraged spatially unsmoothed data—showed nominally significant recruitment during Certain and Uncertain Threat anticipation relative to their respective control conditions (*p*<0.05, uncorrected). For hypothesis testing, we used a standard 2 (*Region:* Ce, BST) × 2 (*Threat*-*Certainty:* Certain, Uncertain) repeated-measures GLM to test potential regional differences in activation during the anticipation of temporally Certain Threat (relative to Certain Safety) versus Uncertain Threat (relative to Uncertain Safety). These analyses leveraged spatially unsmoothed data to maximize anatomical resolution and inferential clarity. Interactions were probed using focal contrasts. Sensitivity analyses confirmed that none of the conclusions materially changed when controlling for potential nuisance variation in mean-centered study, age, and assigned sex (for additional details, see the study OSF collection). A sign test (*Z_Sign_*) was used to nonparametrically test the proportion of participants showing double dissociations.

*Continued…*

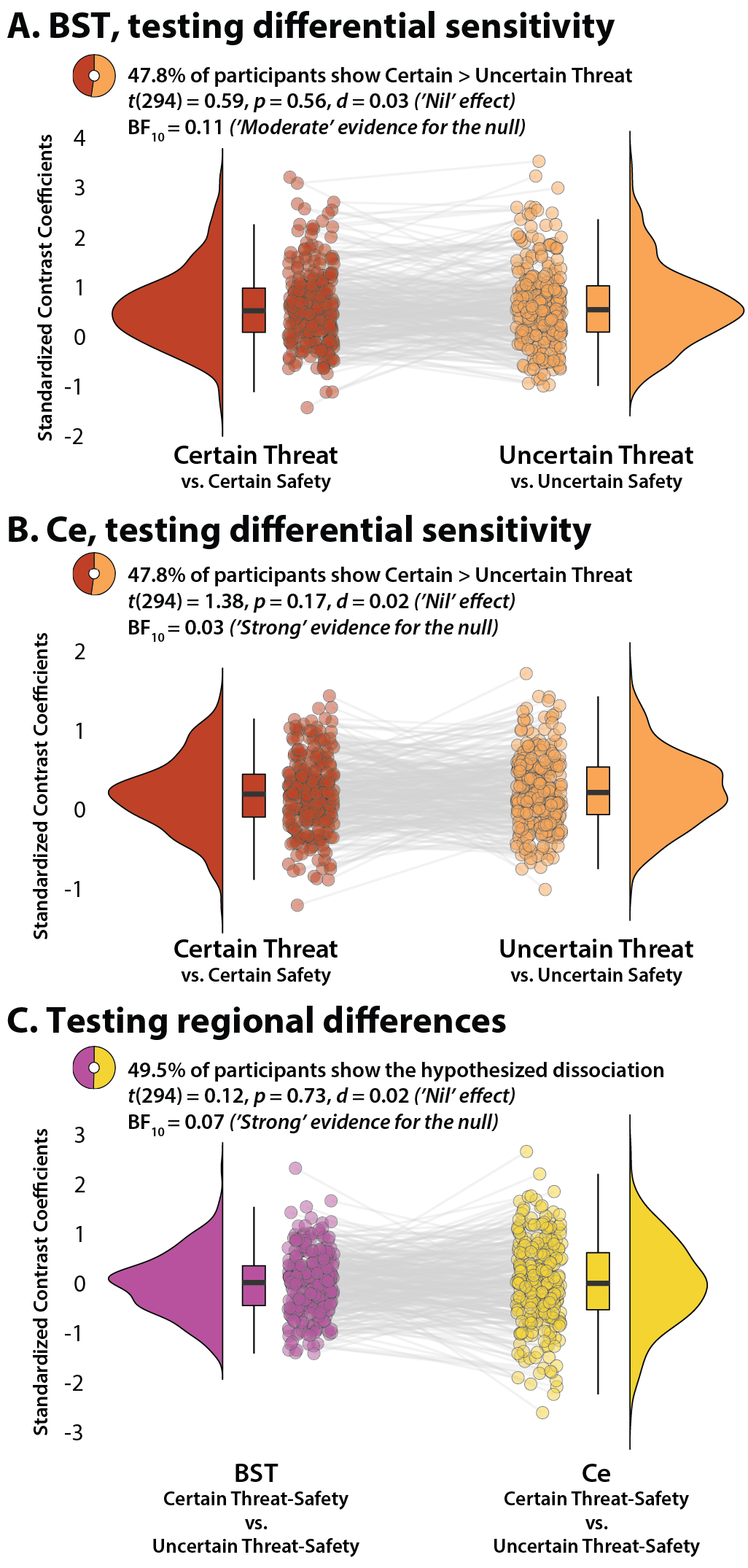
**SUPPLEMENTARY RESULTS**

**Supplementary Figure S2. *The BST and Ce show statistically indistinguishable responses to threat.*** **(A) *BST.*** The raincloud plot depicts BST reactivity to Certain and Uncertain Threat. Ring plot indicates the percentage of participants showing greater BST activation during Certain versus Uncertain threat anticipation. **(B) *Ce.*** The raincloud plot depicts BST reactivity to Certain and Uncertain Threat. Ring plot indicates the percentage of participants showing greater Ce activation during Certain versus Uncertain threat anticipation. Raincloud plots depict the median (*horizontal lines*), interquartile range (*boxes*), individual participants (*dots*), and smoothed data distributions (*half violins*) for each contrast. Whiskers indicate 1.5× the interquartile range. Gray lines depict the sign and magnitude of intra-individual mean differences. Abbreviations—BF, Bayes Factor; BST, bed nucleus of the stria terminalis; Ce, central nucleus of the amygdala; *d*, Cohen’s *d*. **(C) *Testing regional differences.*** The raincloud plot depicts the Region × Condition interaction as a 1-*df* contrast (‘difference of differences’). Ring plot indicates the percentage of participants showing the RDoC-hypothesized dissociation of regional reactivity to threat (BST: Certain < Uncertain Threat; Ce: Certain > Uncertain Threat).

**SUPPLEMENTARY TABLES**

**Supplementary Table S1.** Descriptive statistics for clusters and local extrema showing greater activation during the anticipation of Uncertain Threat relative to Uncertain Safety (FWE *p*<0.05, whole-brain corrected, 4-mm smoothing kernel).

|  |  | **mm^3^** | ***t*** | **x** | **y** | **z** |
| --- | --- | --- | --- | --- | --- | --- |
|  | **Cluster 1** | 516,112 |  |  |  |  |
|  | L Frontal Operculum Cortex |  | 21.66 | -40 | 14 | 2 |
|  | L Insular Cortex |  | 20.23 | -32 | 24 | 6 |
|  | L Cingulate Gyrus, anterior division |  | 19.08 | -6 | 10 | 40 |
|  | L Paracingulate Gyrus |  | 18.97 | -8 | 12 | 38 |
|  | L Temporal Occipital Fusiform Cortex |  | 18.48 | -28 | -60 | -18 |
|  | L Brain Stem |  | 16.62 | -2 | -28 | -2 |
|  | L Left Putamen |  | 16.13 | -22 | 8 | -4 |
|  | L Central Opercular Cortex |  | 15.88 | -52 | 4 | 2 |
|  | L Supramarginal Gyrus, anterior division |  | 15.38 | -64 | -32 | 22 |
|  | L Superior Frontal Gyrus |  | 15.32 | -14 | 2 | 70 |
|  | L Cingulate Gyrus, posterior division |  | 15.25 | -10 | -22 | 40 |
|  | L Supramarginal Gyrus, posterior division |  | 15.12 | -58 | -50 | 34 |
|  | L Precentral Gyrus |  | 15.11 | -40 | -4 | 48 |
|  | L Middle Frontal Gyrus |  | 14.70 | -40 | -2 | 58 |
|  | L Thalamus |  | 14.55 | -10 | -6 | 14 |
|  | L Juxtapositional Lobule Cortex |  | 14.53 | -8 | 2 | 52 |
|  | L Caudate |  | 14.39 | -10 | 0 | 10 |
|  | L Parietal Operculum Cortex |  | 12.56 | -62 | -26 | 18 |
|  | L Superior Parietal Lobule |  | 11.84 | -20 | -50 | 64 |
|  | L Precuneus Cortex |  | 11.64 | -10 | -48 | 52 |
|  | L Postcentral Gyrus |  | 11.22 | -58 | -22 | 24 |
|  | L Inferior Frontal Gyrus, pars opercularis |  | 11.12 | -58 | 10 | 18 |
|  | L Middle Temporal Gyrus, temporooccipital part |  | 10.57 | -46 | -60 | 10 |
|  | L Lateral Occipital Cortex, inferior division |  | 10.04 | -54 | -66 | 10 |
|  | L Lingual Gyrus |  | 9.94 | -12 | -86 | -12 |
|  | L Occipital Pole |  | 9.26 | -20 | -96 | 12 |
|  | L Lateral Occipital Cortex, superior division |  | 9.03 | -32 | -60 | 58 |
|  | L Angular Gyrus |  | 8.97 | -48 | -54 | 52 |
|  | L Pallidum |  | 7.60 | -26 | -14 | -2 |
|  | L Planum Polare |  | 7.35 | -42 | -2 | -14 |
|  | L Temporal Fusiform Cortex, posterior division |  | 6.50 | -44 | -40 | -22 |
|  | L Temporal Pole |  | 5.87 | -46 | 18 | -22 |
|  | R Paracingulate Gyrus |  | 20.98 | 8 | 20 | 36 |
|  | R Frontal Orbital Cortex |  | 20.70 | 34 | 28 | 2 |
|  | R Frontal Operculum Cortex |  | 19.68 | 36 | 22 | 8 |
|  | R Central Opercular Cortex |  | 18.92 | 48 | 6 | 0 |
|  | R Juxtapositional Lobule Cortex |  | 18.84 | 6 | 6 | 48 |
|  | R Putamen |  | 18.23 | 22 | 6 | -6 |
|  | R Cingulate Gyrus, anterior division |  | 18.09 | 10 | 6 | 42 |
|  | R Precentral Gyrus |  | 18.04 | 44 | -2 | 48 |
|  | R Supramarginal Gyrus, posterior division |  | 17.55 | 62 | -42 | 22 |
|  | R Brain Stem |  | 17.44 | 4 | -30 | -2 |
|  | R Superior Frontal Gyrus |  | 17.16 | 2 | 10 | 56 |
|  | R Caudate |  | 17.06 | 12 | -2 | 14 |
|  | R Inferior Frontal Gyrus, pars triangularis |  | 16.12 | 56 | 22 | 0 |
|  | R Cingulate Gyrus, posterior division |  | 15.85 | 10 | -22 | 42 |
|  | R Parietal Operculum Cortex |  | 15.71 | 56 | -28 | 26 |
|  | R Frontal Pole |  | 15.62 | 32 | 48 | 28 |
|  | R Middle Frontal Gyrus |  | 14.95 | 50 | 8 | 42 |
|  | R Temporal Occipital Fusiform Cortex |  | 14.92 | 32 | -54 | -20 |
|  | R Superior Parietal Lobule |  | 14.40 | 24 | -46 | 66 |
|  | R Middle Temporal Gyrus, posterior division |  | 13.51 | 50 | -24 | -6 |
|  | R Angular Gyrus |  | 12.85 | 58 | -50 | 42 |
|  | R Thalamus |  | 12.81 | 6 | -20 | -2 |
|  | R Postcentral Gyrus |  | 12.03 | 32 | -38 | 62 |
|  | R Inferior Frontal Gyrus, pars opercularis |  | 11.81 | 40 | 16 | 26 |
|  | R Precuneus Cortex |  | 11.27 | 8 | -54 | 56 |
|  | R Middle Temporal Gyrus, temporooccipital part |  | 11.23 | 54 | -60 | 6 |
|  | R Occipital Pole |  | 11.13 | 22 | -96 | 16 |
|  | R Lingual Gyrus |  | 9.26 | 10 | -80 | -6 |
|  | R Planum Polare |  | 9.15 | 42 | -4 | -12 |
|  | R Amygdala |  | 9.14 | 28 | -10 | -12 |
|  | R Occipital Fusiform Gyrus |  | 8.91 | 22 | -80 | -12 |
|  | R Lateral Occipital Cortex, inferior division |  | 8.83 | 42 | -78 | -10 |
|  | R Lateral Occipital Cortex, superior division |  | 8.58 | 16 | -80 | 40 |
|  | R Insular Cortex |  | 8.33 | 40 | -16 | -2 |
|  | R Pallidum |  | 7.30 | 14 | -4 | -4 |
|  | R Inferior Temporal Gyrus, temporooccipital part |  | 7.25 | 48 | -54 | -14 |
|  | R Heschls Gyrus (includes H1 and H2) |  | 6.53 | 44 | -12 | 4 |
|  | **Cluster 2** | 15,144 |  |  |  |  |
|  | L Frontal Pole |  | 16.21 | -28 | 48 | 26 |
|  | L Middle Frontal Gyrus |  | 14.75 | -34 | 36 | 40 |
|  | **Cluster 3** | 1,672 |  |  |  |  |
|  | R Temporal Pole |  | 6.08 | 46 | 12 | -42 |
|  | **Cluster 4** | 688 |  |  |  |  |
|  | L Middle Temporal Gyrus, posterior division |  | 6.27 | -60 | -28 | -2 |
|  | **Cluster 5** | 376 |  |  |  |  |
|  | R Frontal Pole |  | 7.49 | 26 | 50 | -14 |
|  | **Cluster 6** | 320 |  |  |  |  |
|  | L Temporal Pole |  | 6.89 | -40 | 6 | -38 |
|  | **Cluster 7** | 272 |  |  |  |  |
|  | L Middle Temporal Gyrus, anterior division |  | 6.50 | -50 | -2 | -30 |
|  | **Cluster 8** | 216 |  |  |  |  |
|  | R Occipital Pole |  | 6.74 | 6 | -96 | -8 |
|  | **Cluster 9** | 176 |  |  |  |  |
|  | R Temporal Pole |  | 6.72 | 52 | 8 | -20 |
|  | **Cluster 10** | 144 |  |  |  |  |
|  | R Parahippocampal Gyrus, anterior division |  | 6.19 | 24 | -8 | -32 |
|  | **Cluster 11** | 104 |  |  |  |  |
|  | L Amygdala |  | 7.67 | -26 | -12 | -12 |
|  | **Cluster 12** | 80 |  |  |  |  |
|  | L Frontal Pole |  | 6.38 | -30 | 54 | -14 |
|  | **Cluster 13** | 64 |  |  |  |  |
|  | L Parahippocampal Gyrus, anterior division |  | 5.89 | -24 | -6 | -32 |
|  | **Cluster 14** | 24 |  |  |  |  |
|  | L Temporal Fusiform Cortex, anterior division |  | 5.22 | -36 | -2 | -42 |
|  | **Cluster 15** | 24 |  |  |  |  |
|  | L Frontal Pole |  | 5.67 | -26 | 46 | -14 |
|  | **Cluster 16** | 24 |  |  |  |  |
|  | R Frontal Pole |  | 5.55 | 30 | 66 | 0 |
|  | **Cluster 17** | 16 |  |  |  |  |
|  | L Parahippocampal Gyrus, anterior division |  | 5.58 | -24 | -10 | -38 |
|  | **Cluster 18** | 16 |  |  |  |  |
|  | R Amygdala |  | 5.53 | 18 | -14 | -12 |
|  | **Cluster 19** | 8 |  |  |  |  |
|  | L Temporal Pole |  | 5.24 | -20 | 2 | -40 |
|  | **Cluster 20** | 8 |  |  |  |  |
|  | R Brain Stem |  | 5.70 | 8 | -24 | -34 |
|  | **Cluster 21** | 8 |  |  |  |  |
|  | R Hippocampus |  | 5.44 | 12 | -12 | -20 |
|  | **Cluster 22** | 8 |  |  |  |  |
|  | L Frontal Pole |  | 5.14 | -24 | 50 | -14 |
|  | **Cluster 23** | 8 |  |  |  |  |
|  | R Frontal Pole |  | 5.13 | 22 | 62 | -8 |
|  | **Cluster 24** | 8 |  |  |  |  |
|  | R Hippocampus |  | 5.15 | 34 | -28 | -6 |
|  | **Cluster 25** | 8 |  |  |  |  |
|  | L Frontal Pole |  | 5.34 | -26 | 66 | -4 |
|  | **Cluster 26** | 8 |  |  |  |  |
|  | L Hippocampus |  | 5.20 | -20 | -38 | -2 |
|  | **Cluster 27** | 8 |  |  |  |  |
|  | R Frontal Pole |  | 5.36 | 14 | 64 | 26 |
|  | **Cluster 28** | 8 |  |  |  |  |
|  | R Postcentral Gyrus |  | 5.42 | 22 | -30 | 62 |

Note: Suprathreshold activation was also evident in the bilateral periaqueductal gray, bilateral bed nucleus of the stria terminalis, and bilateral dorsal amygdala.

**Supplementary Table S2.** Descriptive statistics for clusters and local extrema showing greater activation during the anticipation of Uncertain Safety relative to Uncertain Threat (FWE *p*<0.05, whole-brain corrected, 4-mm smoothing kernel).

|  |  | **mm^3^** | ***t*** | **x** | **y** | **z** |
| --- | --- | --- | --- | --- | --- | --- |
|  | **Cluster 1** | 30,088 |  |  |  |  |
|  | L Intracalcarine Cortex |  | 19.11 | -14 | -80 | 6 |
|  | L Cingulate Gyrus, posterior division |  | 10.74 | -4 | -50 | 14 |
|  | L Lingual Gyrus |  | 10.55 | -10 | -60 | 4 |
|  | L Precuneus Cortex |  | 10.40 | -10 | -62 | 18 |
|  | R Intracalcarine Cortex |  | 18.87 | 14 | -78 | 10 |
|  | R Precuneus Cortex |  | 14.57 | 2 | -68 | 22 |
|  | R Supracalcarine Cortex |  | 13.91 | 2 | -66 | 18 |
|  | R Cingulate Gyrus, posterior division |  | 10.39 | 10 | -50 | 4 |
|  | **Cluster 2** | 5,000 |  |  |  |  |
|  | L Postcentral Gyrus |  | 11.41 | -52 | -10 | 26 |
|  | **Cluster 3** | 3,640 |  |  |  |  |
|  | L Precentral Gyrus |  | 8.43 | -6 | -24 | 60 |
|  | L Postcentral Gyrus |  | 6.85 | -10 | -40 | 70 |
|  | R Precentral Gyrus |  | 10.41 | 2 | -32 | 66 |
|  | **Cluster 4** | 3,112 |  |  |  |  |
|  | R Precentral Gyrus |  | 11.07 | 54 | -6 | 26 |
|  | R Postcentral Gyrus |  | 9.93 | 64 | -6 | 34 |
|  | **Cluster 5** | 2,600 |  |  |  |  |
|  | L Frontal Medial Cortex |  | 5.73 | -6 | 54 | -12 |
|  | R Frontal Pole |  | 9.58 | 0 | 64 | -10 |
|  | R Frontal Medial Cortex |  | 8.36 | 2 | 52 | -12 |
|  | R Subcallosal Cortex |  | 5.49 | 4 | 28 | -18 |
|  | **Cluster 6** | 1,568 |  |  |  |  |
|  | L Lateral Occipital Cortex, superior division |  | 7.85 | -38 | -84 | 38 |
|  | **Cluster 7** | 1,328 |  |  |  |  |
|  | R Lateral Occipital Cortex, superior division |  | 7.61 | 50 | -72 | 36 |
|  | **Cluster 8** | 1,016 |  |  |  |  |
|  | L Insular Cortex |  | 10.02 | -38 | -12 | 14 |
|  | L Central Opercular Cortex |  | 7.88 | -44 | -12 | 18 |
|  | **Cluster 9** | 872 |  |  |  |  |
|  | R Insular Cortex |  | 10.33 | 38 | -8 | 12 |
|  | R Central Opercular Cortex |  | 7.57 | 46 | -10 | 16 |
|  | **Cluster 10** | 576 |  |  |  |  |
|  | R Parahippocampal Gyrus, posterior division |  | 7.93 | 28 | -32 | -18 |
|  | R Lingual Gyrus |  | 6.21 | 26 | -40 | -10 |
|  | **Cluster 11** | 568 |  |  |  |  |
|  | L Parahippocampal Gyrus, posterior division |  | 7.43 | -28 | -38 | -14 |
|  | L Temporal Fusiform Cortex, posterior division |  | 6.31 | -32 | -32 | -18 |
|  | **Cluster 12** | 328 |  |  |  |  |
|  | R Precentral Gyrus |  | 6.19 | 14 | -28 | 66 |
|  | **Cluster 13** | 248 |  |  |  |  |
|  | R Hippocampus |  | 7.49 | 22 | -18 | -20 |
|  | R Amygdala |  | 5.51 | 18 | -8 | -18 |
|  | **Cluster 14** | 184 |  |  |  |  |
|  | L Hippocampus |  | 7.83 | -18 | -16 | -22 |
|  | **Cluster 15** | 112 |  |  |  |  |
|  | R Postcentral Gyrus |  | 6.93 | 12 | -40 | 70 |
|  | **Cluster 16** | 88 |  |  |  |  |
|  | L Cingulate Gyrus, posterior division |  | 5.62 | -6 | -38 | 34 |
|  | **Cluster 17** | 88 |  |  |  |  |
|  | L Precentral Gyrus |  | 6.61 | -12 | -28 | 72 |
|  | **Cluster 18** | 80 |  |  |  |  |
|  | R Frontal Pole |  | 5.91 | 38 | 36 | -12 |
|  | **Cluster 19** | 56 |  |  |  |  |
|  | L Middle Temporal Gyrus, anterior division |  | 5.94 | -64 | -4 | -18 |
|  | **Cluster 20** | 56 |  |  |  |  |
|  | R Superior Frontal Gyrus |  | 5.32 | 20 | 30 | 46 |
|  | **Cluster 21** | 24 |  |  |  |  |
|  | L Hippocampus |  | 5.48 | -22 | -20 | -16 |
|  | **Cluster 22** | 24 |  |  |  |  |
|  | L Frontal Orbital Cortex |  | 5.59 | -34 | 36 | -10 |
|  | **Cluster 23** | 24 |  |  |  |  |
|  | R Lingual Gyrus |  | 5.38 | 18 | -46 | -8 |
|  | **Cluster 24** | 24 |  |  |  |  |
|  | R Subcallosal Cortex |  | 5.79 | 2 | 14 | -6 |
|  | **Cluster 25** | 16 |  |  |  |  |
|  | R Middle Temporal Gyrus, anterior division |  | 5.55 | 64 | 0 | -20 |
|  | **Cluster 26** | 16 |  |  |  |  |
|  | L Precentral Gyrus |  | 5.18 | -12 | -26 | 78 |
|  | **Cluster 27** | 8 |  |  |  |  |
|  | R Subcallosal Cortex |  | 5.21 | 2 | 22 | -24 |
|  | **Cluster 28** | 8 |  |  |  |  |
|  | R Subcallosal Cortex |  | 5.19 | 0 | 26 | -24 |
|  | **Cluster 29** | 8 |  |  |  |  |
|  | L Amygdala |  | 5.32 | -30 | 2 | -18 |
|  | **Cluster 30** | 8 |  |  |  |  |
|  | L Hippocampus |  | 5.38 | -28 | -18 | -16 |
|  | **Cluster 31** | 8 |  |  |  |  |
|  | L Frontal Orbital Cortex |  | 5.11 | -40 | 36 | -14 |
|  | **Cluster 32** | 8 |  |  |  |  |
|  | L Frontal Medial Cortex |  | 5.11 | -6 | 48 | -14 |
|  | **Cluster 33** | 8 |  |  |  |  |
|  | L Frontal Medial Cortex |  | 5.19 | -8 | 52 | -10 |
|  | **Cluster 34** | 8 |  |  |  |  |
|  | R Hippocampus |  | 5.72 | 24 | -26 | -8 |
|  | **Cluster 35** | 8 |  |  |  |  |
|  | R Cuneal Cortex |  | 5.12 | 0 | -86 | 34 |

**Supplementary Table S3.** Descriptive statistics for clusters and local extrema showing greater activation during the anticipation of Certain Threat relative to Certain Safety (FWE *p*<0.05, whole-brain corrected).

|  | | **mm^3^** | ***t*** | **x** | **y** | **z** |
| --- | --- | --- | --- | --- | --- | --- |
|  | **Cluster 1** | 448,752 |  |  |  |  |
|  | L Superior Frontal Gyrus |  | 15.05 | -14 | -2 | 68 |
|  | L Frontal Operculum Cortex |  | 14.25 | -34 | 16 | 10 |
|  | L Putamen |  | 13.61 | -22 | 14 | -6 |
|  | L Cingulate Gyrus, anterior division |  | 13.34 | -8 | 18 | 34 |
|  | L Juxtapositional Lobule Cortex |  | 13.24 | -4 | 4 | 56 |
|  | L Thalamus |  | 13.19 | -10 | -6 | 14 |
|  | L Bed Nucleus of the Stria Terminalis |  | 12.97 | -6 | 4 | 0 |
|  | L Supramarginal Gyrus, posterior division |  | 12.80 | -58 | -50 | 40 |
|  | L Paracingulate Gyrus |  | 12.72 | -8 | 22 | 32 |
|  | L Lateral Occipital Cortex, superior division |  | 12.56 | -36 | -58 | 58 |
|  | L Middle Frontal Gyrus |  | 12.05 | -34 | -4 | 64 |
|  | L Insular Cortex |  | 11.78 | -32 | 26 | 4 |
|  | L Superior Parietal Lobule |  | 11.68 | -34 | -58 | 52 |
|  | L Occipital Fusiform Gyrus |  | 11.61 | -30 | -72 | -20 |
|  | L Brain Stem |  | 11.52 | -2 | -28 | -4 |
|  | L Frontal Pole |  | 11.51 | -30 | 52 | 28 |
|  | L Lingual Gyrus |  | 11.30 | -4 | -74 | -12 |
|  | L Central Opercular Cortex |  | 11.20 | -44 | 4 | 2 |
|  | L Caudate |  | 10.65 | -14 | 18 | -4 |
|  | L Precentral Gyrus |  | 10.62 | -40 | -2 | 46 |
|  | L Frontal Orbital Cortex |  | 10.50 | -32 | 26 | -6 |
|  | L Inferior Frontal Gyrus, pars triangularis |  | 10.16 | -44 | 22 | 8 |
|  | L Angular Gyrus |  | 9.90 | -50 | -56 | 48 |
|  | L Parietal Operculum Cortex |  | 9.77 | -60 | -38 | 24 |
|  | L Precuneus Cortex |  | 9.49 | -10 | -78 | 36 |
|  | L Cingulate Gyrus, posterior division |  | 9.31 | -2 | -26 | 28 |
|  | L Supramarginal Gyrus, anterior division |  | 8.34 | -60 | -38 | 46 |
|  | L Postcentral Gyrus |  | 8.30 | -20 | -32 | 76 |
|  | L Accumbens |  | 7.92 | -8 | 12 | -6 |
|  | L Inferior Frontal Gyrus, pars opercularis |  | 7.69 | -54 | 10 | 16 |
|  | L Middle Temporal Gyrus, temporooccipital part |  | 7.01 | -62 | -56 | 6 |
|  | L Lateral Occipital Cortex, inferior division |  | 6.25 | -56 | -64 | 10 |
|  | L Temporal Pole |  | 6.12 | -50 | 18 | -18 |
|  | L Hippocampus |  | 5.64 | -30 | -12 | -16 |
|  | L Periaqueductal Gray |  | 5.52 | -2 | -34 | -12 |
|  | L Superior Temporal Gyrus, posterior division |  | 5.30 | -66 | -38 | 6 |
|  | R Putamen |  | 15.93 | 22 | 10 | -8 |
|  | R Superior Frontal Gyrus |  | 15.64 | 20 | -8 | 70 |
|  | R Paracingulate Gyrus |  | 15.16 | 8 | 20 | 36 |
|  | R Caudate |  | 14.92 | 12 | -4 | 16 |
|  | R Precentral Gyrus |  | 14.52 | 46 | 0 | 50 |
|  | R Juxtapositional Lobule Cortex |  | 14.08 | 2 | 6 | 48 |
|  | R Bed Nucleus of the Stria Terminalis |  | 13.93 | 8 | 4 | 2 |
|  | R Postcentral Gyrus |  | 13.84 | 34 | -36 | 62 |
|  | R Thalamus |  | 13.41 | 10 | -2 | 12 |
|  | R Frontal Pole |  | 13.02 | 30 | 44 | 24 |
|  | R Superior Parietal Lobule |  | 12.93 | 24 | -44 | 66 |
|  | R Frontal Operculum Cortex |  | 12.55 | 34 | 24 | 8 |
|  | R Supramarginal Gyrus, posterior division |  | 12.54 | 62 | -38 | 32 |
|  | R Middle Frontal Gyrus |  | 12.45 | 38 | -2 | 50 |
|  | R Angular Gyrus |  | 12.29 | 62 | -46 | 30 |
|  | R Central Opercular Cortex |  | 11.69 | 52 | 6 | 2 |
|  | R Lateral Occipital Cortex, superior division |  | 11.46 | 28 | -58 | 62 |
|  | R Brain Stem |  | 10.80 | 4 | -28 | -2 |
|  | R Superior Temporal Gyrus, posterior division |  | 10.67 | 48 | -26 | -4 |
|  | R Cingulate Gyrus, anterior division |  | 10.16 | 0 | 10 | 34 |
|  | R Precuneus Cortex |  | 10.01 | 8 | -56 | 56 |
|  | R Frontal Orbital Cortex |  | 9.63 | 40 | 20 | -8 |
|  | R Cingulate Gyrus, posterior division |  | 9.54 | 10 | -24 | 42 |
|  | R Accumbens |  | 9.49 | 10 | 12 | -4 |
|  | R Parietal Operculum Cortex |  | 9.06 | 56 | -28 | 26 |
|  | R Inferior Frontal Gyrus, pars triangularis |  | 8.66 | 54 | 28 | -6 |
|  | R Supramarginal Gyrus, anterior division |  | 8.36 | 60 | -24 | 28 |
|  | R Middle Temporal Gyrus, posterior division |  | 8.32 | 54 | -22 | -10 |
|  | R Amygdala |  | 8.32 | 30 | -10 | -14 |
|  | R Inferior Frontal Gyrus, pars opercularis |  | 7.95 | 46 | 12 | 28 |
|  | R Lateral Occipital Cortex, inferior division |  | 7.74 | 42 | -84 | -8 |
|  | R Middle Temporal Gyrus, temporooccipital part |  | 7.32 | 60 | -60 | 6 |
|  | R Insular Cortex |  | 6.99 | 40 | 10 | -8 |
|  | R Planum Polare |  | 6.26 | 42 | -4 | -14 |
|  | R Occipital Pole |  | 5.88 | 26 | -94 | 18 |
|  | R Pallidum |  | 5.47 | 16 | -4 | -4 |
|  | R Periaqueductal Gray |  | 5.40 | 2 | -36 | -12 |
|  | R Temporal Occipital Fusiform Cortex |  | 5.30 | 44 | -54 | -16 |
|  | **Cluster 2** | 792 |  |  |  |  |
|  | L Brain Stem |  | 6.73 | -8 | -38 | -46 |
|  | R Brain Stem |  | 8.04 | 2 | -34 | -48 |
|  | **Cluster 3** | 776 |  |  |  |  |
|  | L Lateral Occipital Cortex, inferior division |  | 7.00 | -40 | -82 | -6 |
|  | L Occipital Pole |  | 5.50 | -36 | -92 | -8 |
|  | **Cluster 4** |  |  |  |  |  |
|  | Middle Temporal Gyrus, posterior division |  | 6.95 | -66 | -24 | -6 |
|  | Superior Temporal Gyrus, posterior division |  | 6.71 | -60 | -26 | -2 |
|  | **Cluster 5** | 224 |  |  |  |  |
|  | R Temporal Pole |  | 6.31 | 42 | 6 | -38 |
|  | R Middle Temporal Gyrus, anterior division |  | 6.29 | 48 | 0 | -32 |
|  | **Cluster 6** | 224 |  |  |  |  |
|  | L Frontal Pole |  | 6.05 | -32 | 48 | -14 |
|  | **Cluster 7** | 168 |  |  |  |  |
|  | L Occipital Pole |  | 5.82 | -22 | -96 | 12 |
|  | **Cluster 8** | 168 |  |  |  |  |
|  | R Precentral Gyrus |  | 6.85 | 6 | -30 | 76 |
|  | **Cluster 9** | 152 |  |  |  |  |
|  | R Frontal Pole |  | 6.34 | 40 | 46 | 6 |
|  | **Cluster 10** | 96 |  |  |  |  |
|  | L Hippocampus |  | 6.17 | -22 | -36 | -4 |
|  | **Cluster 11** |  |  |  |  |  |
|  | L Planum Polare |  | 6.74 | -42 | -4 | -14 |
|  | L Insular Cortex |  | 6.58 | -40 | -2 | -16 |
|  | **Cluster 12** | 88 |  |  |  |  |
|  | R Frontal Pole |  | 6.05 | 26 | 48 | -14 |
|  | **Cluster 13** | 80 |  |  |  |  |
|  | R Brain Stem |  | 5.97 | 18 | -34 | -28 |
|  | **Cluster 14** | 72 |  |  |  |  |
|  | L Lateral Occipital Cortex, inferior division |  | 5.73 | -48 | -64 | 4 |
|  | **Cluster 15** | 72 |  |  |  |  |
|  | L Middle Temporal Gyrus, temporooccipital part |  | 6.13 | -46 | -58 | 10 |
|  | **Cluster 16** | 64 |  |  |  |  |
|  | L Brain Stem |  | 6.46 | -6 | -46 | -56 |
|  | **Cluster 17** | 64 |  |  |  |  |
|  | R Brain Stem |  | 5.58 | 0 | -36 | -38 |
|  | **Cluster 18** | 64 |  |  |  |  |
|  | L Brain Stem |  | 6.95 | -4 | -34 | -24 |
|  | **Cluster 19** | 48 |  |  |  |  |
|  | L Brain Stem |  | 6.79 | -12 | -24 | -20 |
|  | **Cluster 20** | 48 |  |  |  |  |
|  | L Lateral Occipital Cortex, inferior division |  | 5.54 | -50 | -74 | -10 |
|  | **Cluster 21** | 40 |  |  |  |  |
|  | L Brain Stem |  | 5.71 | -8 | -38 | -30 |
|  | **Cluster 22** | 40 |  |  |  |  |
|  | L Brain Stem |  | 5.71 | -8 | -20 | -22 |
|  | **Cluster 23** | 32 |  |  |  |  |
|  | L Brain Stem |  | 6.15 | -4 | -18 | -20 |
|  | **Cluster 24** | 32 |  |  |  |  |
|  | R Inferior Temporal Gyrus, temporooccipital part |  | 5.86 | 60 | -58 | -18 |
|  | **Cluster 25** | 32 |  |  |  |  |
|  | R Inferior Temporal Gyrus, temporooccipital part |  | 5.50 | 60 | -58 | -12 |
|  | **Cluster 26** | 32 |  |  |  |  |
|  | R Lateral Occipital Cortex, inferior division |  | 5.44 | 44 | -72 | -4 |
|  | **Cluster 27** | 32 |  |  |  |  |
|  | R Hippocampus |  | 5.63 | 20 | -36 | 6 |
|  | **Cluster 28** | 32 |  |  |  |  |
|  | R Central Opercular Cortex |  | 5.32 | 48 | -18 | 14 |
|  | **Cluster 29** | 16 |  |  |  |  |
|  | R Hippocampus |  | 5.95 | 16 | -12 | -16 |
|  | **Cluster 30** | 16 |  |  |  |  |
|  | R Cingulate Gyrus, posterior division |  | 5.44 | 4 | -38 | 24 |
|  | **Cluster 31** | 16 |  |  |  |  |
|  | L Middle Frontal Gyrus |  | 5.24 | -50 | 22 | 30 |
|  | **Cluster 32** | 8 |  |  |  |  |
|  | R Brain Stem |  | 5.20 | 8 | -36 | -28 |
|  | **Cluster 33** | 8 |  |  |  |  |
|  | R Brain Stem |  | 5.25 | 6 | -34 | -26 |
|  | **Cluster 34** | 8 |  |  |  |  |
|  | L Middle Temporal Gyrus, posterior division |  | 5.17 | -54 | -30 | -6 |
|  | **Cluster 35** | 8 |  |  |  |  |
|  | L Pallidum |  | 5.25 | -20 | -10 | -4 |
|  | **Cluster 36** | 8 |  |  |  |  |
|  | L Frontal Pole |  | 5.26 | -50 | 42 | 0 |
|  | **Cluster 37** | 8 |  |  |  |  |
|  | R Frontal Pole |  | 5.98 | 30 | 68 | 2 |
|  | **Cluster 38** | 8 |  |  |  |  |
|  | L Lateral Occipital Cortex, inferior division |  | 5.43 | -54 | -70 | 6 |
|  | **Cluster 39** | 8 |  |  |  |  |
|  | L Lateral Occipital Cortex, inferior division |  | 5.52 | -42 | -72 | 10 |
|  | **Cluster 40** | 8 |  |  |  |  |
|  | R Lateral Occipital Cortex, inferior division |  | 6.02 | 42 | -68 | 10 |
|  | **Cluster 41** | 8 |  |  |  |  |
|  | R Lateral Occipital Cortex, inferior division |  | 5.24 | 42 | -62 | 12 |
|  | **Cluster 42** | 8 |  |  |  |  |
|  | R Parietal Operculum Cortex |  | 5.20 | 46 | -22 | 16 |
|  | **Cluster 43** | 8 |  |  |  |  |
|  | R Occipital Pole |  | 5.22 | 18 | -98 | 18 |
|  | **Cluster 44** | 8 |  |  |  |  |
|  | L Angular Gyrus |  | 5.35 | -58 | -60 | 18 |
|  | **Cluster 45** | 8 |  |  |  |  |
|  | R Inferior Frontal Gyrus, pars triangularis |  | 5.40 | 50 | 26 | 18 |
|  | **Cluster 46** | 8 |  |  |  |  |
|  | R Lateral Occipital Cortex, superior division |  | 5.26 | 26 | -84 | 36 |

Note: Suprathreshold activation was also evident in the bilateral dorsal amygdala.

**Supplementary Table S4.** Descriptive statistics for clusters and local extrema showing greater activation during the anticipation of Certain Safety relative to Certain Threat (FWE *p*<0.05, whole-brain corrected, 4-mm smoothing kernel).

|  | | **mm^3^** | ***t*** | **x** | **y** | **z** |
| --- | --- | --- | --- | --- | --- | --- |
|  | **Cluster 1** | 35,472 |  |  |  |  |
|  | L Intracalcarine Cortex |  | 19.18 | -10 | -76 | 10 |
|  | L Lingual Gyrus |  | 10.42 | -20 | -44 | -10 |
|  | L Precuneus Cortex |  | 8.68 | -6 | -60 | 12 |
|  | L Occipital Pole |  | 8.46 | -10 | -98 | -4 |
|  | L Parahippocampal Gyrus, posterior division |  | 7.48 | -30 | -34 | -16 |
|  | L Cingulate Gyrus, posterior division |  | 6.42 | -4 | -48 | 14 |
|  | R Intracalcarine Cortex |  | 20.15 | 14 | -82 | 4 |
|  | R Lingual Gyrus |  | 17.44 | 0 | -74 | 6 |
|  | R Occipital Pole |  | 15.03 | 6 | -90 | 6 |
|  | R Precuneus Cortex |  | 13.09 | 0 | -68 | 18 |
|  | R Parahippocampal Gyrus, posterior division |  | 8.09 | 20 | -34 | -16 |
|  | R Occipital Fusiform Gyrus |  | 8.01 | 28 | -68 | -6 |
|  | R Cingulate Gyrus, posterior division |  | 7.98 | 10 | -50 | 6 |
|  | R Temporal Fusiform Cortex, posterior division |  | 7.67 | 30 | -34 | -18 |
|  | **Cluster 2** | 288 |  |  |  |  |
|  | L Postcentral Gyrus |  | 6.49 | -62 | -8 | 22 |
|  | **Cluster 3** | 272 |  |  |  |  |
|  | L Subcallosal Cortex |  | 6.25 | -2 | 16 | -6 |
|  | R Subcallosal Cortex |  | 7.09 | 2 | 14 | -10 |
|  | **Cluster 4** | 168 |  |  |  |  |
|  | R Frontal Medial Cortex |  | 5.70 | 2 | 44 | -14 |
|  | **Cluster 5** | 160 |  |  |  |  |
|  | L Lingual Gyrus |  | 6.47 | -28 | -58 | -6 |
|  | **Cluster 6** | 40 |  |  |  |  |
|  | R Frontal Medial Cortex |  | 5.48 | 0 | 50 | -14 |
|  | **Cluster 7** | 32 |  |  |  |  |
|  | L Temporal Pole |  | 5.58 | -42 | 12 | -18 |
|  | **Cluster 8** | 32 |  |  |  |  |
|  | R Subcallosal Cortex |  | 5.56 | 0 | 30 | -16 |
|  | **Cluster 9** | 32 |  |  |  |  |
|  | R Precentral Gyrus |  | 5.34 | 56 | -6 | 24 |
|  | **Cluster 10** | 24 |  |  |  |  |
|  | R Parahippocampal Gyrus, anterior division |  | 5.94 | 20 | -20 | -22 |
|  | **Cluster 11** | 16 |  |  |  |  |
|  | L Frontal Medial Cortex |  | 5.54 | -2 | 36 | -18 |
|  | **Cluster 12** | 16 |  |  |  |  |
|  | R Right Hippocampus |  | 6.65 | 24 | -26 | -8 |
|  | **Cluster 13** | 16 |  |  |  |  |
|  | R Precentral Gyrus |  | 5.39 | 60 | -4 | 28 |
|  | **Cluster 14** | 8 |  |  |  |  |
|  | L Middle Temporal Gyrus, anterior division |  | 5.29 | -56 | -6 | -16 |
|  | **Cluster 15** | 8 |  |  |  |  |
|  | L Frontal Medial Cortex |  | 5.16 | -8 | 42 | -14 |
|  | **Cluster 16** | 8 |  |  |  |  |
|  | L Frontal Pole |  | 5.15 | -4 | 56 | -12 |
|  | **Cluster 17** | 8 |  |  |  |  |
|  | R Postcentral Gyrus |  | 5.44 | 66 | -6 | 22 |

**Supplementary Table S5.** Descriptive statistics for clusters and local extrema showing greater activation during the anticipation of Uncertain Threat relative to Certain Threat (FWE *p*<0.05, whole-brain corrected).

|  |  | **mm^3^** | ***t*** | **x** | **y** | **z** |
| --- | --- | --- | --- | --- | --- | --- |
|  | **Cluster 1** | 142,448 |  |  |  |  |
|  | L Occipital Pole |  | 13.58 | -12 | -94 | -8 |
|  | L Temporal Occipital Fusiform Cortex |  | 11.63 | -24 | -54 | -16 |
|  | L Occipital Fusiform Gyrus |  | 11.36 | -20 | -86 | -10 |
|  | L Cingulate Gyrus, anterior division |  | 9.84 | -6 | 8 | 40 |
|  | L Lateral Occipital Cortex, inferior division |  | 9.30 | -32 | -86 | -18 |
|  | L Superior Frontal Gyrus |  | 8.75 | -14 | 10 | 66 |
|  | L Paracingulate Gyrus |  | 8.64 | -6 | 28 | 30 |
|  | L Temporal Fusiform Cortex, posterior division |  | 5.82 | -42 | -40 | -24 |
|  | R Occipital Fusiform Gyrus |  | 15.81 | 30 | -78 | -10 |
|  | R Frontal Orbital Cortex |  | 15.80 | 34 | 28 | 2 |
|  | R Frontal Operculum Cortex |  | 15.11 | 42 | 24 | 4 |
|  | R Occipital Pole |  | 14.00 | 26 | -94 | 12 |
|  | R Precentral Gyrus |  | 13.40 | 42 | -2 | 52 |
|  | R Lateral Occipital Cortex, inferior division |  | 13.31 | 44 | -80 | -10 |
|  | R Central Opercular Cortex |  | 12.41 | 56 | 4 | 4 |
|  | R Temporal Occipital Fusiform Cortex |  | 12.37 | 36 | -46 | -18 |
|  | R Middle Frontal Gyrus |  | 12.06 | 42 | 0 | 60 |
|  | R Inferior Frontal Gyrus, pars triangularis |  | 12.02 | 54 | 22 | 2 |
|  | R Parietal Operculum Cortex |  | 12.00 | 54 | -28 | 26 |
|  | R Juxtapositional Lobule Cortex |  | 11.96 | 4 | 4 | 46 |
|  | R Supramarginal Gyrus, anterior division |  | 11.81 | 60 | -26 | 24 |
|  | R Cingulate Gyrus, anterior division |  | 11.78 | 8 | 6 | 42 |
|  | R Paracingulate Gyrus |  | 11.62 | 2 | 18 | 38 |
|  | R Superior Frontal Gyrus |  | 10.58 | 4 | 20 | 58 |
|  | R Inferior Frontal Gyrus, pars opercularis |  | 10.43 | 60 | 12 | 8 |
|  | R Supramarginal Gyrus, posterior division |  | 10.37 | 58 | -44 | 30 |
|  | R Putamen |  | 10.28 | 28 | 2 | -6 |
|  | R Cingulate Gyrus, posterior division |  | 10.02 | 12 | -22 | 42 |
|  | R Angular Gyrus |  | 9.92 | 62 | -48 | 34 |
|  | R Middle Temporal Gyrus, temporooccipital part |  | 8.46 | 46 | -56 | 6 |
|  | R Inferior Temporal Gyrus, temporooccipital part |  | 6.80 | 46 | -42 | -16 |
|  | R Planum Polare |  | 6.49 | 40 | -18 | -2 |
|  | R Frontal Pole |  | 6.14 | 54 | 38 | -8 |
|  | R Insular Cortex |  | 5.73 | 38 | -14 | 4 |
|  | R Pallidum |  | 5.49 | 16 | 2 | 0 |
|  | R Middle Temporal Gyrus, posterior division |  | 5.36 | 56 | -34 | 0 |
|  | **Cluster 2** | 13,728 |  |  |  |  |
|  | L Insular Cortex |  | 16.18 | -32 | 24 | 6 |
|  | L Frontal Operculum Cortex |  | 14.44 | -44 | 20 | -4 |
|  | L Inferior Frontal Gyrus, pars opercularis |  | 10.78 | -50 | 20 | 4 |
|  | L Central Opercular Cortex |  | 10.68 | -56 | 2 | 4 |
|  | L Precentral Gyrus |  | 8.83 | -56 | 8 | 2 |
|  | **Cluster 3** | 4,056 |  |  |  |  |
|  | L Supramarginal Gyrus, posterior division |  | 8.05 | -60 | -50 | 44 |
|  | L Supramarginal Gyrus, anterior division |  | 8.04 | -64 | -32 | 22 |
|  | L Parietal Operculum Cortex |  | 7.77 | -62 | -30 | 20 |
|  | L Angular Gyrus |  | 7.19 | -60 | -54 | 38 |
|  | **Cluster 4** | 2,272 |  |  |  |  |
|  | R Superior Parietal Lobule |  | 10.02 | 22 | -46 | 66 |
|  | R Postcentral Gyrus |  | 6.80 | 34 | -38 | 62 |
|  | **Cluster 5** | 1,280 |  |  |  |  |
|  | L Frontal Pole |  | 7.03 | -38 | 46 | 30 |
|  | **Cluster 6** | 1,232 |  |  |  |  |
|  | L Brain Stem |  | 6.84 | -2 | -30 | -2 |
|  | R Brain Stem |  | 9.53 | 4 | -30 | -2 |
|  | R Thalamus |  | 7.70 | 6 | -22 | -2 |
|  | **Cluster 7** | 1,032 |  |  |  |  |
|  | R Thalamus |  | 7.98 | 10 | 0 | 8 |
|  | R Caudate |  | 6.20 | 16 | -10 | 20 |
|  | **Cluster 8** | 776 |  |  |  |  |
|  | R Frontal Pole |  | 6.68 | 32 | 50 | 26 |
|  | **Cluster 9** | 744 |  |  |  |  |
|  | L Putamen |  | 8.43 | -26 | 6 | -8 |
|  | L Insular Cortex |  | 6.78 | -36 | -2 | -6 |
|  | **Cluster 10** | 680 |  |  |  |  |
|  | L Precentral Gyrus |  | 7.07 | -36 | -4 | 48 |
|  | L Middle Frontal Gyrus |  | 6.44 | -38 | -2 | 56 |
|  | **Cluster 11** | 608 |  |  |  |  |
|  | R Middle Temporal Gyrus, posterior division |  | 7.23 | 52 | -22 | -6 |
|  | **Cluster 12** | 552 |  |  |  |  |
|  | L Cingulate Gyrus, posterior division |  | 6.42 | -2 | -16 | 28 |
|  | R Cingulate Gyrus, posterior division |  | 7.41 | 4 | -26 | 26 |
|  | **Cluster 13** | 216 |  |  |  |  |
|  | L Brain Stem |  | 7.06 | -6 | -36 | -48 |
|  | **Cluster 14** | 200 |  |  |  |  |
|  | L Superior Parietal Lobule |  | 6.31 | -18 | -52 | 64 |
|  | **Cluster 15** | 176 |  |  |  |  |
|  | L Cingulate Gyrus, posterior division |  | 7.15 | -12 | -24 | 40 |
|  | **Cluster 16** | 104 |  |  |  |  |
|  | L Brain Stem |  | 6.27 | -8 | -28 | -18 |
|  | **Cluster 17** | 104 |  |  |  |  |
|  | L Inferior Temporal Gyrus, temporooccipital part |  | 5.82 | -44 | -60 | -12 |
|  | **Cluster 18** | 72 |  |  |  |  |
|  | L Thalamus |  | 5.82 | -6 | -14 | -2 |
|  | **Cluster 19** | 40 |  |  |  |  |
|  | L Brain Stem |  | 6.29 | -6 | -32 | -8 |
|  | **Cluster 20** | 40 |  |  |  |  |
|  | L Lateral Occipital Cortex, superior division |  | 5.70 | -30 | -82 | 18 |
|  | **Cluster 21** | 24 |  |  |  |  |
|  | L Brain Stem |  | 5.77 | -4 | -36 | -30 |
|  | **Cluster 22** | 24 |  |  |  |  |
|  | L Middle Frontal Gyrus |  | 5.30 | -44 | 34 | 36 |
|  | **Cluster 23** | 16 |  |  |  |  |
|  | R Heschls Gyrus (includes H1 and H2) |  | 5.34 | 46 | -14 | 6 |
|  | **Cluster 24** | 16 |  |  |  |  |
|  | L Inferior Frontal Gyrus, pars opercularis |  | 5.19 | -56 | 12 | 18 |
|  | **Cluster 25** | 16 |  |  |  |  |
|  | R Cingulate Gyrus, anterior division |  | 5.37 | 8 | 38 | 18 |
|  | **Cluster 26** | 16 |  |  |  |  |
|  | R Lateral Occipital Cortex, superior division |  | 5.22 | 34 | -72 | 24 |
|  | **Cluster 27** | 8 |  |  |  |  |
|  | L Temporal Pole |  | 5.23 | -42 | 6 | -42 |
|  | **Cluster 28** | 8 |  |  |  |  |
|  | L Planum Polare |  | 5.51 | -40 | -20 | -4 |
|  | **Cluster 29** | 8 |  |  |  |  |
|  | R Insular Cortex |  | 5.20 | 36 | -22 | 6 |
|  | **Cluster 30** | 8 |  |  |  |  |
|  | R Angular Gyrus |  | 5.17 | 48 | -54 | 56 |

**Supplementary Table S6.** Descriptive statistics for clusters and local extrema showing greater activation during the anticipation of Certain Threat relative to Uncertain Threat (FWE *p*<0.05, whole-brain corrected, 4-mm smoothing kernel).

|  |  | **mm^3^** | ***t*** | **x** | **y** | **z** |
| --- | --- | --- | --- | --- | --- | --- |
|  | **Cluster 1** | 35,560 |  |  |  |  |
|  | L Precuneus Cortex |  | 11.99 | -12 | -62 | 18 |
|  | L Lingual Gyrus |  | 11.19 | -12 | -54 | 0 |
|  | L Cingulate Gyrus, posterior division |  | 8.89 | -8 | -44 | 4 |
|  | L Hippocampus |  | 7.12 | -20 | -38 | 2 |
|  | L Thalamus |  | 6.87 | -14 | -36 | 4 |
|  | R Precuneus Cortex |  | 14.01 | 14 | -58 | 16 |
|  | R Cingulate Gyrus, posterior division |  | 11.01 | 10 | -48 | 4 |
|  | R Cuneal Cortex |  | 10.29 | 4 | -72 | 28 |
|  | R Lateral Occipital Cortex, superior division |  | 10.12 | 48 | -72 | 34 |
|  | R Hippocampus |  | 8.10 | 18 | -36 | 4 |
|  | **Cluster 2** | 21,344 |  |  |  |  |
|  | L Precentral Gyrus |  | 10.80 | -56 | -8 | 46 |
|  | L Postcentral Gyrus |  | 9.92 | -10 | -42 | 64 |
|  | R Postcentral Gyrus |  | 10.73 | 2 | -34 | 66 |
|  | R Precentral Gyrus |  | 8.82 | 6 | -22 | 66 |
|  | **Cluster 3** | 5,208 |  |  |  |  |
|  | L Lateral Occipital Cortex, superior division |  | 9.77 | -38 | -80 | 36 |
|  | **Cluster 4** | 4,400 |  |  |  |  |
|  | R Postcentral Gyrus |  | 9.34 | 66 | -10 | 30 |
|  | R Precentral Gyrus |  | 8.55 | 60 | -6 | 42 |
|  | **Cluster 5** | 2,104 |  |  |  |  |
|  | L Frontal Medial Cortex |  | 6.63 | -4 | 52 | -8 |
|  | R Frontal Pole |  | 9.06 | 0 | 64 | -10 |
|  | R Frontal Medial Cortex |  | 7.28 | 2 | 50 | -12 |
|  | **Cluster 6** | 1,904 |  |  |  |  |
|  | R Hippocampus |  | 11.81 | 22 | -20 | -18 |
|  | **Cluster 7** | 1,448 |  |  |  |  |
|  | L Hippocampus |  | 10.29 | -22 | -20 | -16 |
|  | L Amygdala |  | 5.86 | -12 | -6 | -20 |
|  | **Cluster 8** | 1,168 |  |  |  |  |
|  | L Central Opercular Cortex |  | 9.57 | -38 | -12 | 18 |
|  | L Insular Cortex |  | 7.60 | -40 | -8 | 6 |
|  | **Cluster 9** | 1,048 |  |  |  |  |
|  | L Parahippocampal Gyrus, posterior division |  | 7.98 | -24 | -38 | -16 |
|  | L Temporal Fusiform Cortex, posterior division |  | 7.86 | -34 | -34 | -18 |
|  | **Cluster 10** | 856 |  |  |  |  |
|  | R Lingual Gyrus |  | 9.44 | 26 | -38 | -12 |
|  | R Parahippocampal Gyrus, posterior division |  | 8.43 | 26 | -34 | -16 |
|  | R Temporal Fusiform Cortex, posterior division |  | 8.13 | 34 | -36 | -14 |
|  | **Cluster 11** | 656 |  |  |  |  |
|  | R Superior Frontal Gyrus |  | 7.01 | 22 | 30 | 44 |
|  | **Cluster 12** | 576 |  |  |  |  |
|  | R Insular Cortex |  | 9.14 | 38 | -8 | 12 |
|  | R Central Opercular Cortex |  | 5.85 | 46 | -10 | 16 |
|  | **Cluster 13** | 536 |  |  |  |  |
|  | R Middle Temporal Gyrus, anterior division |  | 6.69 | 60 | -4 | -16 |
|  | **Cluster 14** | 376 |  |  |  |  |
|  | R Subcallosal Cortex |  | 8.48 | 2 | 12 | -4 |
|  | **Cluster 15** | 272 |  |  |  |  |
|  | R Lateral Occipital Cortex, superior division |  | 6.03 | 20 | -66 | 50 |
|  | **Cluster 16** | 136 |  |  |  |  |
|  | L Juxtapositional Lobule Cortex |  | 7.66 | -4 | 0 | 64 |
|  | **Cluster 17** | 120 |  |  |  |  |
|  | R Postcentral Gyrus |  | 6.09 | 48 | -16 | 52 |
|  | **Cluster 18** | 112 |  |  |  |  |
|  | L Middle Temporal Gyrus, temporooccipital part |  | 6.07 | -60 | -62 | -8 |
|  | L Lateral Occipital Cortex, inferior division |  | 5.57 | -58 | -68 | -8 |
|  | **Cluster 19** | 96 |  |  |  |  |
|  | L Superior Frontal Gyrus |  | 5.67 | -22 | 26 | 46 |
|  | **Cluster 20** | 88 |  |  |  |  |
|  | L Precuneus Cortex |  | 5.52 | -2 | -60 | 64 |
|  | **Cluster 21** | 64 |  |  |  |  |
|  | R Frontal Pole |  | 6.26 | 0 | 62 | 6 |
|  | **Cluster 22** | 48 |  |  |  |  |
|  | R Frontal Pole |  | 5.50 | 10 | 70 | 12 |
|  | **Cluster 23** | 48 |  |  |  |  |
|  | L Lateral Occipital Cortex, superior division |  | 5.76 | -8 | -86 | 42 |
|  | **Cluster 24** | 40 |  |  |  |  |
|  | L Lateral Occipital Cortex, superior division |  | 5.42 | -16 | -84 | 48 |
|  | **Cluster 25** | 32 |  |  |  |  |
|  | R Accumbens |  | 6.01 | 12 | 14 | -8 |
|  | **Cluster 26** | 32 |  |  |  |  |
|  | R Caudate |  | 6.12 | 6 | 10 | -2 |
|  | **Cluster 27** | 32 |  |  |  |  |
|  | R Precuneus Cortex |  | 5.50 | 12 | -50 | 42 |
|  | **Cluster 28** | 24 |  |  |  |  |
|  | L Middle Temporal Gyrus, anterior division |  | 5.12 | -62 | -4 | -16 |
|  | **Cluster 29** | 24 |  |  |  |  |
|  | L Frontal Pole |  | 5.78 | -20 | 66 | 12 |
|  | **Cluster 30** | 24 |  |  |  |  |
|  | L Planum Temporale |  | 5.37 | -48 | -38 | 16 |
|  | **Cluster 31** | 24 |  |  |  |  |
|  | L Supramarginal Gyrus, posterior division |  | 5.38 | -56 | -42 | 18 |
|  | **Cluster 32** | 24 |  |  |  |  |
|  | L Frontal Pole |  | 5.65 | -10 | 70 | 18 |
|  | **Cluster 33** | 24 |  |  |  |  |
|  | L Superior Parietal Lobule |  | 5.20 | -28 | -54 | 52 |
|  | **Cluster 34** | 16 |  |  |  |  |
|  | R Frontal Medial Cortex |  | 5.29 | 2 | 40 | -18 |
|  | **Cluster 35** | 16 |  |  |  |  |
|  | L Lateral Occipital Cortex, superior division |  | 5.17 | -6 | -72 | 58 |
|  | **Cluster 36** | 8 |  |  |  |  |
|  | L Amygdala |  | 5.31 | -30 | 2 | -18 |
|  | **Cluster 37** | 8 |  |  |  |  |
|  | R Frontal Pole |  | 5.17 | 36 | 38 | -8 |
|  | **Cluster 38** | 8 |  |  |  |  |
|  | L Lateral Occipital Cortex, superior division |  | 5.28 | -30 | -76 | 48 |
|  | **Cluster 39** | 8 |  |  |  |  |
|  | R Lateral Occipital Cortex, superior division |  | 5.37 | 20 | -64 | 60 |

**SUPPLEMENTARY REFERENCES**

Anderson, J. S., Druzgal, T. J., Lopez-Larson, M., Jeong, E. K., Desai, K., & Yurgelun-Todd, D. (2011). Network anticorrelations, global regression, and phase-shifted soft tissue correction. *Hum Brain Mapp*, *32*(6), 919-934. <https://doi.org/10.1002/hbm.21079>

Avants, B. B., Tustison, N. J., Song, G., Cook, P. A., Klein, A., & Gee, J. C. (2011). A reproducible evaluation of ANTs similarity metric performance in brain image registration [Article]. *Neuroimage*, *54*, 2033-2044. <https://doi.org/10.1016/j.neuroimage.2010.09.025>

Avery, S. N., Clauss, J. A., & Blackford, J. U. (2016). The human BNST: Functional role in anxiety and addiction. *Neuropsychopharmacology*, *41*, 126-141. <https://doi.org/10.1038/npp.2015.185>

Bach, D. R., Castegnetti, G., Korn, C. W., Gerster, S., Melinscak, F., & Moser, T. (2018). Psychophysiological modeling: Current state and future directions. *Psychophysiology*, *55*, e13214. <https://doi.org/10.1111/psyp.13209>

Bach, D. R., Flandin, G., Friston, K. J., & Dolan, R. J. (2010). Modelling event-related skin conductance responses. *Int J Psychophysiol*, *75*, 349-356. <https://doi.org/10.1016/j.ijpsycho.2010.01.005>

Bach, D. R., & Friston, K. J. (2013). Model-based analysis of skin conductance responses: Towards causal models in psychophysiology. *Psychophysiology*, *50*, 15-22. <https://doi.org/10.1111/j.1469-8986.2012.01483.x>

BIAC. (2022). *IXI Dataset*. Imperial College London. Retrieved April 19 from <https://brain-development.org/ixi-dataset/>

Bo, K., Kraynak, T. E., Kwon, M., Sun, M., Gianaros, P. J., & Wager, T. D. (2024). A systems identification approach using Bayes factors to deconstruct the brain bases of emotion regulation. *Nat Neurosci*, *27*, 975-987. <https://doi.org/10.1038/s41593-024-01605-7>

Cohen, J. (1988). *Statistical power analysis for the behavioral sciences* (2nd ed.). Lawrence Erlbaum Associates.

Cohen, J. R. (1994). The earth is round (p < .05). *American Psychologist*, *49*, 997-1003. <https://doi.org/10.1037/0003-066X.49.12.997>

Cornwell, B. R., Didier, P. R., Grogans, S. E., Anderson, A. S., Islam, S., Kim, H. C., . . . Shackman, A. J. (2025). A shared threat-anticipation circuit is dynamically engaged at different moments by certain and uncertain threat. *Journal of Neuroscience*, *45*, e2113242025. <https://doi.org/10.1523/JNEUROSCI.2113-24.2025>

Cox, R. W. (1996). AFNI: Software for analysis and visualization of functional magnetic resonance neuroimages. *Computers and Biomedical Research*, *29*, 162-173.

Daldrup, T., Remmes, J., Lesting, J., Gaburro, S., Fendt, M., Meuth, P., . . . Seidenbecher, T. (2015). Expression of freezing and fear-potentiated startle during sustained fear in mice. *Genes Brain Behav*, *14*, 281-291. <https://doi.org/10.1111/gbb.12211>

Daniel-Watanabe, L., & Fletcher, P. C. (2022). Are fear and anxiety truly distinct? *Biological Psychiatry Global Open Science*, *2*, 341-349. <https://doi.org/https://doi.org/10.1016/j.bpsgos.2021.09.006>

Davis, M., Walker, D. L., Miles, L., & Grillon, C. (2010). Phasic vs sustained fear in rats and humans: Role of the extended amygdala in fear vs anxiety. *Neuropsychopharmacology*, *35*, 105-135. <https://doi.org/10.1038/npp.2009.109>

Desikan, R. S., Ségonne, F., Fischl, B., Quinn, B. T., Dickerson, B. C., Blacker, D., . . . Killiany, R. J. (2006). An automated labeling system for subdividing the human cerebral cortex on MRI scans into gyral based regions of interest. *Neuroimage*, *31*, 968-980.

Domschke, K. (2022). Fear and anxiety-Distinct or "kindred" phenomena? *Biol Psychiatry Glob Open Sci*, *2*, 314-315. <https://doi.org/10.1016/j.bpsgos.2022.07.001>

Edlow, B. L., Takahashi, E., Wu, O., Benner, T., Dai, G., Bu, L., . . . Folkerth, R. D. (2012). Neuroanatomic connectivity of the human ascending arousal system critical to consciousness and its disorders. *J Neuropathol Exp Neurol*, *71*(6), 531-546. <https://doi.org/10.1097/NEN.0b013e3182588293>

Eskildsen, S. F., Coupé, P., Fonov, V., Manjón, J. V., Leung, K. K., Guizard, N., . . . Alzheimer's Disease Neuroimaging Initiative. (2012). BEaST: brain extraction based on nonlocal segmentation technique. *Neuroimage*, *59*, 2362-2373.

Fox, A. S., Lapate, R. C., Davidson, R. J., & Shackman, A. J. (2018). The nature of emotion: A research agenda for the 21st century. In A. S. Fox, R. C. Lapate, A. J. Shackman, & R. J. Davidson (Eds.), *The nature of emotion. Fundamental questions* (2nd ed., pp. 403-417). Oxford University Press.

Fox, A. S., & Shackman, A. J. (2019). The central extended amygdala in fear and anxiety: Closing the gap between mechanistic and neuroimaging research. *Neuroscience letters*, *693*, 58-67. <https://doi.org/https://doi.org/10.1016/j.neulet.2017.11.056>

Frazier, J. A., Chiu, S., Breeze, J. L., Makris, N., Lange, N., Kennedy, D. N., . . . Biederman, J. (2005). Structural brain magnetic resonance imaging of limbic and thalamic volumes in pediatric bipolar disorder. *American Journal of Psychiatry*, *162*, 1256-1265.

Grabner, G., Janke, A. L., Budge, M. M., Smith, D., Pruessner, J., & Collins, D. L. (2006). Symmetric atlasing and model based segmentation: an application to the hippocampus in older adults. *Med Image Comput Comput Assist Interv Int Conf Med Image Comput Comput Assist Interv*, *9*, 58–66. <https://doi.org/https://doi.org/10.1007/11866763_8>

Grogans, S. E., Bliss-Moreau, E., Buss, K. A., Clark, L. A., Fox, A. S., Keltner, D., . . . Shackman, A. J. (2023). The nature and neurobiology of fear and anxiety: State of the science and opportunities for accelerating discovery. *Neuroscience & Biobehavioral Reviews*, *151*, 105237. <https://doi.org/https://doi.org/10.1016/j.neubiorev.2023.105237>

Grogans, S. E., Hur, J., Barstead, M. G., Anderson, A. S., Islam, S., Kuhn, M., . . . Shackman, A. J. (2024). Neuroticism/negative emotionality is associated with increased reactivity to uncertain threat in the bed nucleus of the stria terminalis, not the amygdala. *Journal of Neuroscience*, *44*, e1868232024.

Gronau, Q. F., Ly, A., & Wagenmakers, E.-J. (2020). Informed Bayesian t-Tests. *The American Statistician*, *74*, 137-143. <https://doi.org/10.1080/00031305.2018.1562983>

Grupe, D. W., & Nitschke, J. B. (2013). Uncertainty and anticipation in anxiety: an integrated neurobiological and psychological perspective. *Nat Rev Neurosci*, *14*, 488-501. <https://doi.org/nrn3524> [pii]

10.1038/nrn3524

Gungor, N. Z., & Paré, D. (2016). Functional heterogeneity in the bed nucleus of the stria terminalis. *Journal of Neuroscience*, *36*, 8038-8049.

Hefner, K. R., Moberg, C. A., Hachiya, L. Y., & Curtin, J. J. (2013). Alcohol stress response dampening during imminent versus distal, uncertain threat. *J Abnorm Psychol*, *122*, 756-769. <https://doi.org/10.1037/a0033407>

Henson, R. (2007). Efficient experimental design for fMRI. In K. Friston, J. Ashburner, S. Kiebel, T. Nichols, & W. Penny (Eds.), *Statistical Parametric Mapping: The Analysis of Functional Brain Images* (pp. 193-210). Academic Press.

Hur, J., Smith, J. F., DeYoung, K. A., Anderson, A. S., Kuang, J., Kim, H. C., . . . Shackman, A. J. (2020). Anxiety and the neurobiology of temporally uncertain threat anticipation. *Journal of Neuroscience*, *40*, 7949-7964. <https://doi.org/10.1101/2020.02.25.964734>

Jenkinson, M., Beckmann, C. F., Behrens, T. E., Woolrich, M. W., & Smith, S. M. (2012). FSL. *Neuroimage*, *62*, 782-790. <https://doi.org/10.1016/j.neuroimage.2011.09.015>

Kim, H. C., Kaplan, C. M., Islam, S., Anderson, A. S., Piper, M. E., Bradford, D. E., . . . Shackman, A. J. (2023). Acute nicotine abstinence amplifies subjective withdrawal symptoms and threat-evoked fear and anxiety, but not extended amygdala reactivity. *PLoS One*, *18*, e0288544. <https://doi.org/https://doi.org/10.1371/journal.pone.0288544>

Kruger, O., Shiozawa, T., Kreifelts, B., Scheffler, K., & Ethofer, T. (2015). Three distinct fiber pathways of the bed nucleus of the stria terminalis to the amygdala and prefrontal cortex. *Cortex*, *66*, 60-68. <https://doi.org/10.1016/j.cortex.2015.02.007>

Lange, M. D., Daldrup, T., Remmers, F., Szkudlarek, H. J., Lesting, J., Guggenhuber, S., . . . Pape, H. C. (2017). Cannabinoid CB1 receptors in distinct circuits of the extended amygdala determine fear responsiveness to unpredictable threat. *Mol Psychiatry*, *22*, 1422-1430. <https://doi.org/10.1038/mp.2016.156>

Lebow, M. A., & Chen, A. (2016). Overshadowed by the amygdala: the bed nucleus of the stria terminalis emerges as key to psychiatric disorders. *Mol Psychiatry*, *21*, 450-463. <https://doi.org/10.1038/mp.2016.1>

LeDoux, J. E., & Pine, D. S. (2016). Using neuroscience to help understand fear and anxiety: A two-system framework. *Am J Psychiatry*, *173*, 1083-1093. <https://doi.org/10.1176/appi.ajp.2016.16030353>

Lorio, S., Fresard, S., Adaszewski, S., Kherif, F., Chowdhury, R., Frackowiak, R. S., . . . Draganski, B. (2016). New tissue priors for improved automated classification of subcortical brain structures on MRI. *Neuroimage*, *130*, 157-166. <https://doi.org/10.1016/j.neuroimage.2016.01.062>

Love, J., Selker, R., Marsman, M., Jamil, T., Dropmann, D., Verhagen, J., . . . Wagenmakers, E.-J. (2019). JASP: Graphical statistical software for common statistical designs. *Journal of Statistical Software*, *88*, 1-17. <https://doi.org/10.18637/jss.v088.i02>

Makris, N., Goldstein, J. M., Kennedy, D., Hodge, S. M., Caviness, V. S., Faraone, S. V., . . . Seidman, L. J. (2006). Decreased volume of left and total anterior insular lobule in schizophrenia. *Schizophrenia Research*, *83*, 155-171.

McCormick, M., Liu, X., Jomier, J., Marion, C., & Ibanez, L. (2014). ITK: enabling reproducible research and open science. *Frontiers in Neuroinformatics*, *8*, 13. <https://doi.org/10.3389/fninf.2014.00013>

Miles, L., Davis, M., & Walker, D. (2011). Phasic and sustained fear are pharmacologically dissociable in rats. *Neuropsychopharmacology*, *36*, 1563-1574. <https://doi.org/10.1038/npp.2011.29>

Mobbs, D., Headley, D. B., Ding, W., & Dayan, P. (2020). Space, time, and fear: Survival computations along defensive circuits. *Trends in cognitive sciences*, *24*, 228-241. <https://doi.org/10.1016/j.tics.2019.12.016>

Moberg, C. A., Bradford, D. E., Kaye, J. T., & Curtin, J. J. (2017). Increased startle potentiation to unpredictable stressors in alcohol dependence: Possible stress neuroadaptation in humans. *Journal of Abnormal Psychology*, *126*, 441-453. <https://doi.org/10.1037/abn0000265>

Moscarello, J. M., & Penzo, M. A. (2022). The central nucleus of the amygdala and the construction of defensive modes across the threat-imminence continuum. *Nat Neurosci*, *25*, 999-1008. <https://doi.org/10.1038/s41593-022-01130-5>

Nichols, T., Brett, M., Andersson, J., Wager, T., & Poline, J. B. (2005). Valid conjunction inference with the minimum statistic. *Neuroimage*, *25*, 653-660. <https://doi.org/S1053-8119(04)00750-5> [pii]

10.1016/j.neuroimage.2004.12.005 [doi]

NIMH. (2011). *Negative valence systems: Workshop proceedings (March 13, 2011 – March 15, 2011; Rockville, Maryland)*. Retrieved July 1 from <https://www.nimh.nih.gov/research/research-funded-by-nimh/rdoc/negative-valence-systems-workshop-proceedings.shtml>

Orederu, T., Lennon, V., Vervliet, B., & Schiller, D. (2024). Fear. In A. Scarantino (Ed.), *Emotion Theory* (Vol. II, pp. 152-175). Routledge.

Perusini, J. N., & Fanselow, M. S. (2015). Neurobehavioral perspectives on the distinction between fear and anxiety. *Learn Mem*, *22*, 417-425. <https://doi.org/10.1101/lm.039180.115>

Poldrack, R. A., Baker, C. I., Durnez, J., Gorgolewski, K. J., Matthews, P. M., Munafo, M. R., . . . Yarkoni, T. (2017). Scanning the horizon: towards transparent and reproducible neuroimaging research. *Nat Rev Neurosci*, *18*, 115-126. <https://doi.org/10.1038/nrn.2016.167>

Pruim, R. H. R., Mennes, M., van Rooij, D., Llera, A., Buitelaar, J. K., & Beckmann, C. F. (2015). ICA-AROMA: a robust ICA-based strategy for removing motion artifacts from fMRI data. *Neuroimage*, *112*, 267-277.

Rorden, C. (2019, September 2, 2019). *MRIcron*. NITRC. Retrieved April 18 from <https://www.nitrc.org/projects/mricron>

Schimmack, U. (2019). *Statistics wars: Don’t change alpha. Change the null-hypothesis!* Retrieved December 15 from <https://replicationindex.com/category/nil-hypothesis/>

Schmalz, X., Biurrun Manresa, J., & Zhang, L. (2023). What is a Bayes factor? *Psychological Methods*, *28*, 705-718. <https://doi.org/10.1037/met0000421>

Schönbrodt, F. D., Wagenmakers, E. J., Zehetleitner, M., & Perugini, M. (2017). Sequential hypothesis testing with Bayes factors: Efficiently testing mean differences. *Psychol Methods*, *22*, 322-339. <https://doi.org/10.1037/met0000061>

Shackman, A. J., & Fox, A. S. (2016). Contributions of the central extended amygdala to fear and anxiety. *Journal of Neuroscience*, *36*, 8050-8063. <https://doi.org/10.1523/JNEUROSCI.0982-16.2016>

Shackman, A. J., Tromp, D. P. M., Stockbridge, M. D., Kaplan, C. M., Tillman, R. M., & Fox, A. S. (2016). Dispositional negativity: An integrative psychological and neurobiological perspective. *Psychological Bulletin*, *142*, 1275-1314.

ten Donkelaar, H. J., Tzourio-Mazoyer, N., & Mai, J. K. (2018). Toward a common terminology for the gyri and sulci of the human cerebral cortex [Review]. *Frontiers in Neuroanatomy*, *12*, 93. <https://doi.org/10.3389/fnana.2018.00093>

Theiss, J. D., Ridgewell, C., McHugo, M., Heckers, S., & Blackford, J. U. (2017). Manual segmentation of the human bed nucleus of the stria terminalis using 3T MRI. *Neuroimage*, *146*, 288-292. <https://doi.org/10.1016/j.neuroimage.2016.11.047>

Tillman, R. M., Stockbridge, M. D., Nacewicz, B. M., Torrisi, S., Fox, A. S., Smith, J. F., & Shackman, A. J. (2018). Intrinsic functional connectivity of the central extended amygdala. *Human Brain Mapping*, *39*, 1291-1312.

Tovote, P., Fadok, J. P., & Lüthi, A. (2015). Neuronal circuits for fear and anxiety. *Nat Rev Neurosci*, *16*, 317-331. <https://doi.org/10.1038/nrn3945>

Tukey, J. W. (1977). *Exploratory data analysis*. Addison Wesley.

Tustison, N. J., Avants, B. B., Cook, P. A., Zheng, Y. J., Egan, A., Yushkevich, P. A., & Gee, J. C. (2010). N4ITK: Improved N3 bias correction [Article]. *IEEE Transactions on Medical Imaging*, *29*, 1310-1320. <https://doi.org/10.1109/tmi.2010.2046908>

van Doorn, J., van den Bergh, D., Böhm, U., Dablander, F., Derks, K., Draws, T., . . . Wagenmakers, E.-J. (2021). The JASP guidelines for conducting and reporting a Bayesian analysis. *Psychonomic Bulletin & Review*, *28*, 813-826. <https://doi.org/10.3758/s13423-020-01798-5>

Wagenmakers, E.-J., Love, J., Marsman, M., Jamil, T., Ly, A., Verhagen, J., . . . Morey, R. D. (2018). Bayesian inference for psychology. Part II: Example applications with JASP. *Psychonomic Bulletin & Review*, *25*, 58-76. <https://doi.org/10.3758/s13423-017-1323-7>

Walter, A., Mai, J., Lanta, L., & Görcs, T. (1991). Differential distribution of immunohistochemical markers in the bed nucleus of the stria terminalis in the human brain. *J Chem Neuroanat*, *4*(4), 281-298.

Wellcome Centre for Human Neuroimaging. (2022). *SPM*. University College London. Retrieved April 18 from <https://fil.ion.ucl.ac.uk/spm/>

Wickham, H. (2016). *ggplot2: Elegant graphics for data analysis* (2nd ed.). Springer-Verlag.
