## Supplementary figures and images for "Fear, anxiety, and the extended amygdala—*Absence of evidence for strict functional segregation*"

### Supp Fig 1

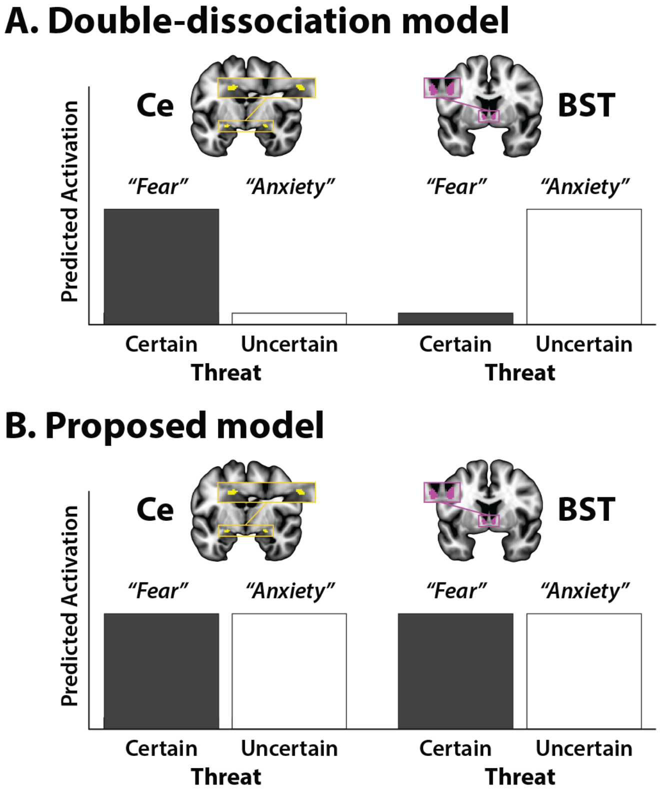

### Supp Fig 2

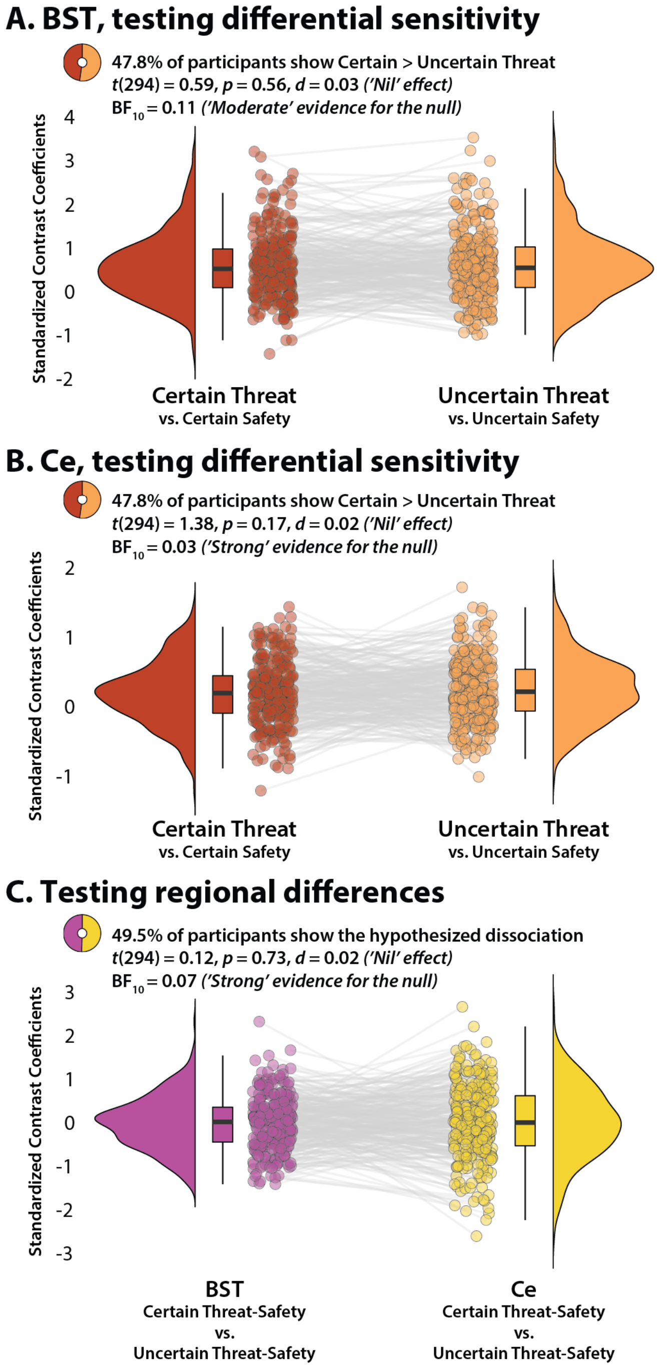
